# Stabilizing mammalian RNA thermometer confers neuroprotection in subarachnoid hemorrhage

**DOI:** 10.1101/2024.05.17.594628

**Authors:** Min Zhang, Bin Zhang, Chengli Liu, Marco Preußner, Megha Ayachit, Weiming Li, Yafei Huang, Deyi Liu, Quanwei He, Ann-Kathrin Emmerichs, Stefan Meinke, Shu Chen, Lin Wang, Liduan Zheng, Qiubo Li, Qin Huang, Tom Haltenhof, Ruoxi Gao, Xianan Qin, Aifang Cheng, Tianzi Wei, Li Yu, Mario Schubert, Xin Gao, Mingchang Li, Florian Heyd

## Abstract

Mammals tightly regulate their core body temperature, yet how cells sense and respond to small temperature changes at the molecular level remains incompletely understood. Here, we discover a significant enrichment of RNA G-quadruplex (rG4) motifs around splice sites of cold-repressed exons. These thermosensing RNA structures, when stabilized, mask splice sites, reducing exon inclusion. Focusing on cold-induced neuroprotective RBM3, we demonstrate that rG4s near splice sites of a cold-repressed poison exon are stabilized at low temperatures, leading to exon exclusion. This enables evasion of nonsense-mediated decay, increasing RBM3 expression at cold. Additionally, increasing intracellular potassium concentration stabilizes rG4s and enhances RBM3 expression, leading to RBM3-dependent neuroprotection in a mouse model of subarachnoid hemorrhage. Our findings unveil a mechanism how mammalian RNAs directly sense temperature and potassium perturbations, integrating them into gene expression programs. This opens new avenues for treating diseases arising from splicing defects and disorders benefiting from therapeutic hypothermia.

**Highlights:**

1. rG4s are enriched near splice sites of cassette exons repressed upon cold shock
2. rG4s act as RNA thermometers in mammals by controlling accessibility of splice sites
3. rG4 stability mediates temperature-dependent RBM3 expression *ex vivo* and *in vivo*
4. Stabilizing rG4s in RBM3 exon 3a reduces brain damage in subarachnoid hemorrhage

**Graphical abstract:** 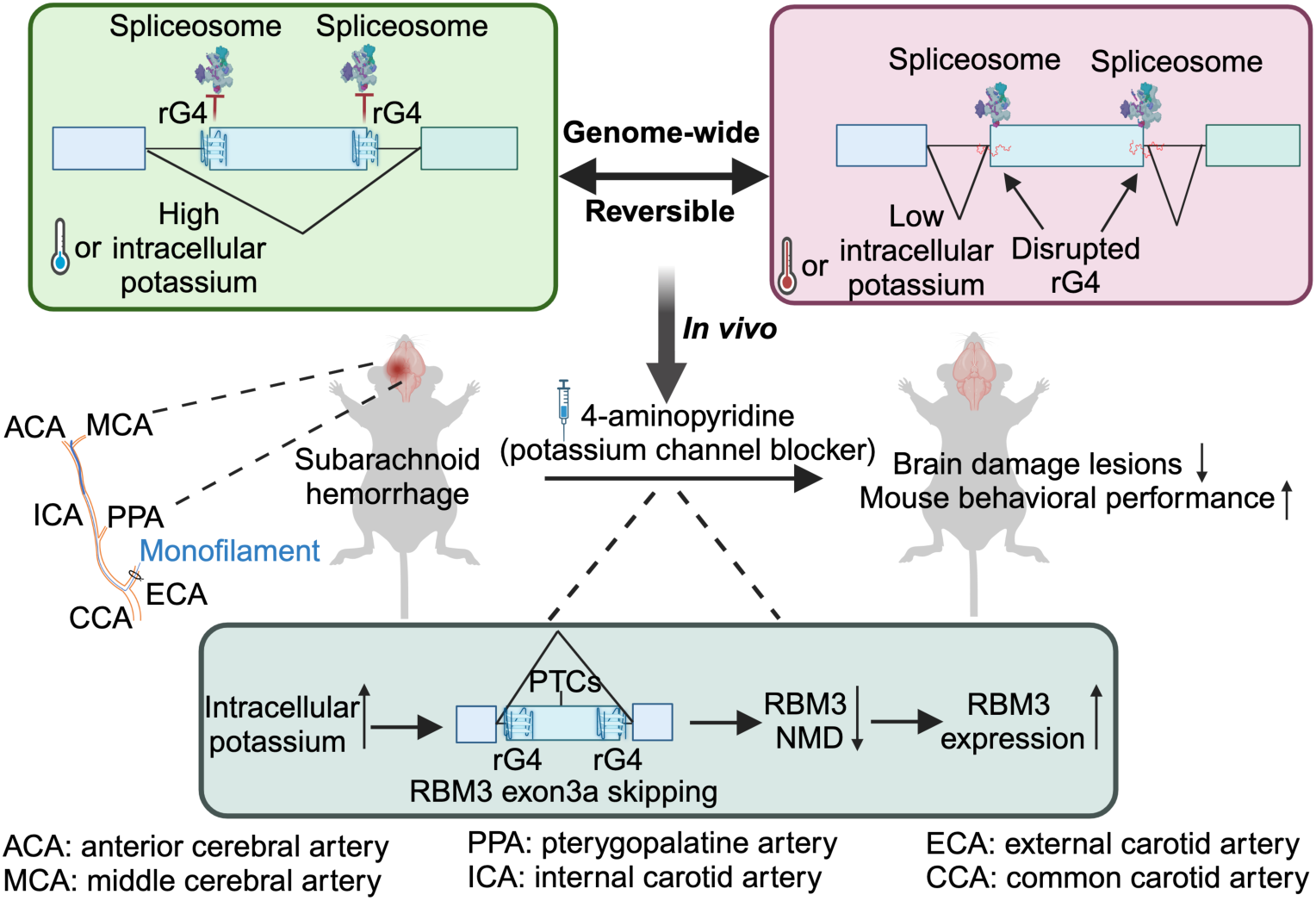

Here is a refined model of RNA G-quadruplexes acting as evolutionarily conserved thermo- and potassium sensors, modulating alternative splicing in mammals. RNA G-quadruplexes (rG4s) function as reversible temperature sensors, impacting alternative splicing dynamics. In low temperatures or high potassium conditions, rG4s can mask surrounding splice sites, rendering these sites inaccessible, thereby promoting exon skipping. Conversely, at high temperatures or under low potassium conditions, rG4s become destabilized, allowing splice sites to be exposed, and facilitating efficient exon inclusion. RBM3, a well-known cold-induced protein with neuroprotective functions, harbors a poison exon with rG4s around splice sites, that, upon inclusion, triggers NMD (non-sense mediated decay) of the RBM3 mRNA. Under low temperatures or high potassium conditions, rG4s shield the splice sites, leading to poison exon skipping and increased RBM3 expression. Stabilization of these rG4s through increased K^+^ promotes poison exon skipping, enabling escape from NMD, and ultimately elevating RBM3 expression. Notably, 4-AP, a clinically used pan voltage-gated potassium channel blocker, protects against neuronal damage in a subarachnoid hemorrhage mouse model in an RBM-dependent manner. (ISS: intronic splicing silencer, ESE: exon splicing enhancer, ACA: anterior cerebral artery, MCA: middle cerebral artery, PPA: pterygopalatine artery, ICA: internal carotid artery, ECA: external carotid artery, CCA: common carotid artery)

## Introduction

In most mammals, including humans, core body temperature is rigorously regulated to stay around ∼37°C^1,2^, with subtle circadian fluctuations within a range of ∼1-4°C^3^. Consequently, cells in most organs experience a relatively narrow temperature spectrum. Certain organs, however, maintain distinct temperatures, for example lower temperature in skin and testis, temperature gradients in the reproductive tract^4–6^ and higher temperature in brain^7^. Significant deviations from normothermia lead to heightened morbidity and mortality^8,9^. Accidental hypothermia, characterized by a prolonged reduction in body temperature below 35°C profoundly disrupts the normal functioning of physiological systems^10–12^. Conversely, extreme hyperthermia, culminating in cell death and heatstroke, occurs when the core body temperature exceeds 40.5°C^13^. Therapeutic hypothermia (TH) has long been recognized as a neuroprotective strategy, with numerous recent clinical trials exploring its potential for neuronal protection. However, the therapeutic window for hypothermia is limited to 1 to 3 days due to associated side effects. The primary latent phase of cellular injury following brain damage lasts approximately 6 hours, followed by a secondary phase of deterioration extending over several days^14^. Despite its promise, the underlying molecular mechanisms of TH and strategies to ensure its continuous application for neuronal protection remain unclear.

At the molecular level, numerous studies have investigated cellular responses to temperature fluctuations using heat-shock and cold-shock treatments. The well-established understanding is that transient receptor potential (TRP) channels expressed in primary sensory neurons directly perceive temperature changes^15–18^. Additionally, even minor temperature shifts of 1°C, suffice to influence RNA splicing by altering the phosphorylation status of SR proteins^19,20^. In our prior work, we demonstrated that this regulatory process relies on the activity of CDC-like kinase (CLK) and is governed by conformational rearrangements within the kinase’s active center at different temperatures^21^. Beyond the temperature-dependent regulation through altered channel or kinase activity, RNA itself may function as a sensor for temperature variations due to its structural flexibility, which arises from alternative base pairing and interactions. Such RNA thermometers have been well studied in bacteria^22^, yeasts^23^ and plants^24–26^. However, the role of RNA secondary structures as direct thermo-sensors remains unexplored in mammals^27,28^.

G-quadruplexes (G4s) represent comparably stable secondary structures formed by Hoogsteen guanine base pairs (G-G). Notably, RNA G-rich elements, which were suggested to form rG4 *in vitro* and in cells^29^, play crucial roles in diverse biological processes, contributing to multiple molecular functions^30,31^. In humans, RNA G-rich sequences have been identified as enhancers of alternative polyadenylation in LDL Receptor Related Protein 5 (LRP5) and FMR1 Autosomal Homolog 1 (FXR1), leading to the generation of shorter transcripts^32^. They also function as common neurite localization signal to transport the mRNA of post-synaptic density proteins within neurites^33^, contribute to the phase separation of RNA granules^34,35^, and regulate translation^36,37^. Furthermore, RNA G-rich sequences may repress the recognition of 5’-splice sites by recruiting heterogeneous nuclear ribonucleoprotein H1 (hnRNP H1)^38,39^, but may also increase splicing efficiency^40,41^, for example through recruiting heterogeneous nuclear ribonucleoprotein F (hnRNP F)^42^, which may act through destabilizing G-quadruplexes^43,44^. Furthermore, a recent genome-wide analysis in mammalian cells found an enrichment of rG4s near splice sites and suggests that rG4s could facilitate exon inclusion by recruiting RNA-binding proteins^45^. However, the involvement of rG4s as dynamic, temperature-dependent regulatory RNA structures in mammals has not been investigated.

Here, we show a substantial enrichment of RNA G-quadruplex motifs near splice sites of cassette exons whose inclusion is repressed at lower temperatures in mammals. We show that splicing changes of cassette exons following cold shock exhibit a global negative correlation with rG4 scores. Using the neuroprotective and cold-induced RBM3 as an example, we characterize a specific rG4 in detail. We find that stabilizing rG4s located proximal to the splice sites of RBM3’s cold-repressed poison exon^46,47^, either through low temperature or high potassium concentration, promotes exon skipping, thereby increasing RBM3 expression by evading nonsense-mediated decay (NMD)^46,47^. Importantly, increasing intracellular potassium level via a potassium channel blocker (4-aminopyridine, 4-AP) stabilizes rG4s and confers RBM3-dependent neuroprotection in a subarachnoid hemorrhage mouse model. Our findings suggest a novel neuroprotective mechanism for 4-AP, involving the stabilization of a mammalian RNA thermometer that regulates alternative splicing coupled to NMD, thereby controlling RBM3 expression, with potential applications in protecting neurons in diverse human diseases.

## Results

### rG4 motifs are significantly enriched around splice sites of cold-repressed cassette exons

Our previous research has illuminated the role of kinases, specifically CLK1/4, in governing temperature-dependent alternative splicing within mammalian cells. In this mechanism, CLKs modulate alternative splicing by affecting the phosphorylation of SR proteins^21^. Amongst others, we performed RNA sequencing in HEK293T cells at different temperatures (35°C and 39°C) that were treated with TG003 (a CLK-specific inhibitor)^21^ or okadaic acid (OA, a broad phosphatase inhibitor), with DMSO as control (**Figure S1A)**. We observed that pharmacological perturbation of phosphorylation via TG003 or okadaic had a stronger influence on splicing at high temperatures than at low temperatures (**Figures 1A** and **S1B)**. This suggests that phosphorylation-dependent splicing regulation has a stronger impact at high temperatures, while another mechanism, not or less dependent on phosphorylation, preferentially impacts splicing at low temperatures. Furthermore, while around 60% (2,453 out of 3,981) of temperature-dependent splicing events responded to CLK inhibition, a substantial proportion of exons (∼40%) was not impacted^21^. This points to the presence of other molecular mechanisms modulating temperature-dependent alternative splicing.

**Figure 1.**
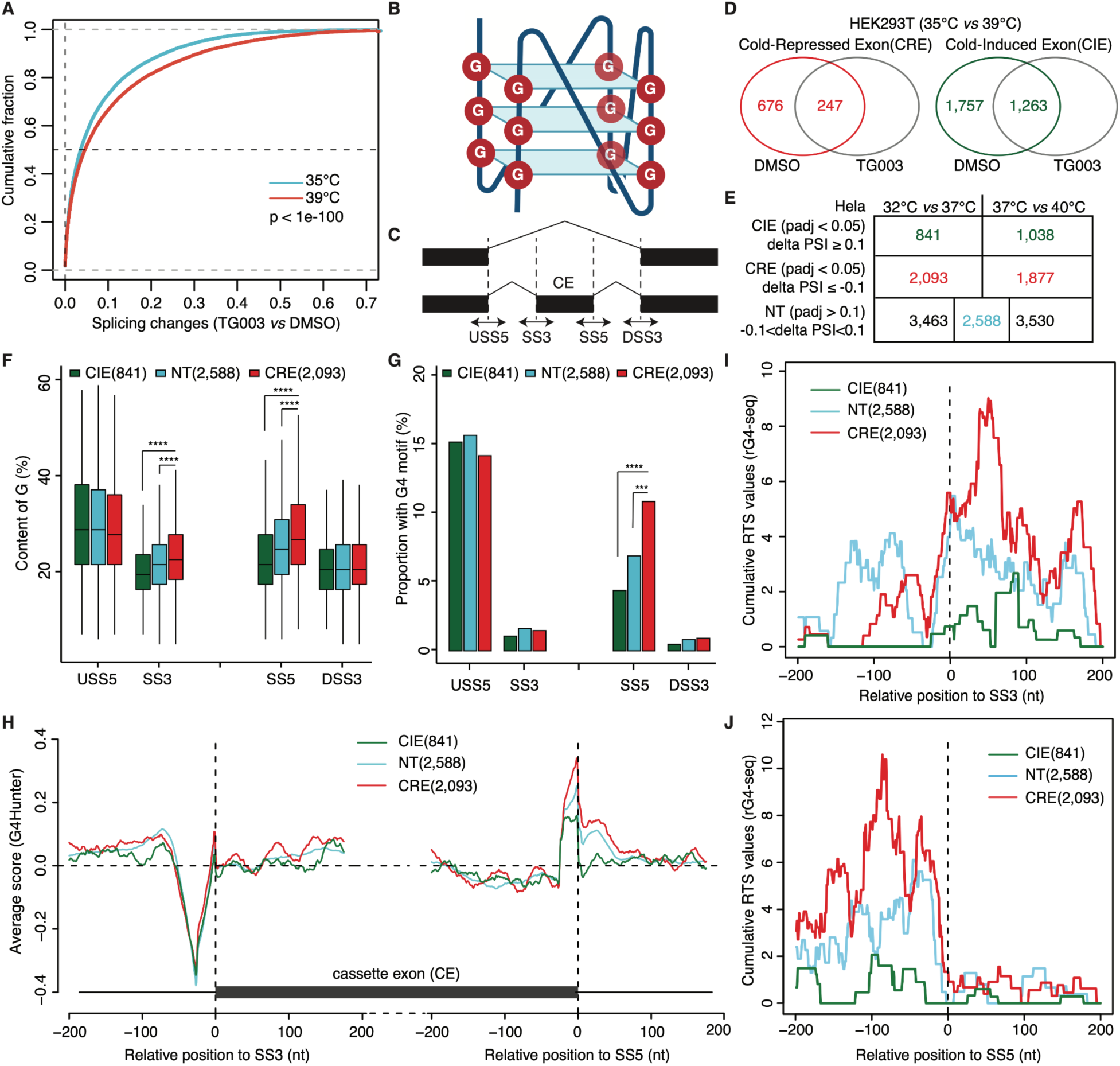
G4 motifs are enriched around splice sites of cold-repressed exons (CREs). (A) TG003 treatment exerting a more pronounced influence on splicing changes at high temperature (39°C) compared to low temperature (35°C), as demonstrated by the cumulative curves of absolute delta PSI values (TG003 *vs* DMSO) in HEK293T cells. (B) Schematic representation of a parallel G4 structure. (C) Schematic depicting the analyzed sequence in four regions around splice sites (SS) of each cassette exon (CE). (D) Venn diagram showing the number of cold-repressed exons (CREs) and cold-induced exons (CIEs) in HEK293T cells treated with DMSO or TG003 at 35°C *vs* 39°C (also see Methods). (E) The number of CIEs, CREs and non-temperature-sensitive exons (NTs) in Hela cells in two comparisons (32°C *vs* 37°C, 37°C *vs* 40°C). The final NT exons for further analysis were selected as the intersections of NT in these two comparisons. (F) Distribution of G-content in sequences within four regions (see C) of CREs, CIEs and NTs in Hela cells (32°C *vs* 37°C). Significance was estimated by Wilcox test and only the significant results compared to CRE were indicated in the figure. ****, p < 0.0001. (G) Proportion of exons with G4 motifs within four regions (see C) of CREs, CIEs and NTs in Hela cells (32°C *vs* 37°C). Significance was estimated by hypergeometric test and only the significant results compared to CRE were indicated in the figure. ****, p < 0.0001. ***, p < 0.001. (H) The average G4 scores predicted by G4Hunter at each nucleotide from −200 to 200 around splice sites in CREs, CIEs and NTs. (I) and (J) Cumulative Reverse Transcriptase Stalling (RTS) values based on rG4-seq^50^ at the 3’ splice site (SS3, I) or the 5’ splice site (SS5, J) of in CREs, CIEs and NTs.

It has been reported that RNA secondary structures serve as molecular thermometers in bacteria^22^, yeasts^22^, and plants^24,25^. For example, rG4 formation in plants is globally enhanced in response to cold (4°C) temperature to increase RNA stability^24^. Hence, we reasoned that rG4 structures **(Figure 1B**) would be well-suited to act as thermo-sensors in mammalian cells. To inspect this possibility, we calculated the G content for sequences surrounding splice sites of cassette exons, using upstream and downstream constitutive exons as controls (**Figure 1C**). We classified the cassette exons into three categories, including cold-repressed exons (CREs), non-temperature sensitive exons (NTs) (**Figure S1C**) and cold-induced exons (CIEs) based on their percent spliced-in (PSI) at 35°C and 39°C. To explore the potential association between rG4-based regulation and kinase-based regulation, we defined CLK-dependent temperature-sensitive exons by requiring that CREs or CIEs are specifically changing in DMSO but not in TG003 treated samples, as well as CLK-independent ones using the overlapped CREs or CIEs between DMSO and TG003 treated samples (**Figure 1D**). We found that in both CLK-independent and CLK-dependent temperature-sensitive exons, G-content of sequences flanking the splice sites of CREs was significantly higher than that in CIEs and NTs. Such differences were not observed for sequences flanking splice sites of upstream and downstream constitutive exons, making a higher G-content specific for cold-repressed cassette exons (**Figures S1D** and **S1E)**. As this holds true also for CREs that are independent of CLK activity, these G-rich sequences may contribute to an alternative mechanism controlling temperature-sensitive alternative splicing.

To further validate these findings in a different cell culture model, we performed RNA-Seq in Hela cells at 32°C, 37°C and 40°C (**Figure S1F)**. Within these RNA-Seq data, we identified 2,093 CREs and 841 CIEs upon cold shock (32°C *vs* 37°C) **(Figure 1E)**. Consistently, we observed a significantly higher G content in sequences flanking the splice sites of CREs compared with that of CIEs and NTs (**Figure 1F**). However, for those exons with splicing changes upon heat shock (40°C *vs* 37°C), the enrichment was not observed (**Figure S1G**), suggesting G-rich sequences may preferentially function in lower temperatures. Next, we used G4 motifs from a published dataset^48^ and searched for putative rG4s in the vicinity of splice sites of cold-induced cassette exons at 32°C, as well as around the 5’-splice sites of the upstream and the 3’-splice sites of the downstream flanking exon as controls. Indeed, we observed a strong enrichment of rG4 motifs in sequences flanking the 5’-splice sites of CREs compared to that from NTs and CIEs (**Figure 1G**). It is worth noting that rG4 motifs were also present close to the 5’-splice sites of the upstream constitutive exons, which has been previously reported and may be related to a splicing-enhancing function of rG4s^45^. To confirm the presence of rG4s close to 5’-splice sites of CREs and to address the position of rG4s with respect to splice sites at single nucleotide resolution, we scanned sequences flanking each cassette exon with a 25 nt window using G4Hunter^49^. In line with a previous study^45^, the highest G-quadruplex (G4) scores are found around the 5’-splice sites of cassette exons. More importantly, the scores were higher for CREs than for CIEs and NTs, with the G4 score of 5’-splice sites of cold-induced exons being close to 0 (**Figure 1H**). To further validate our *in silico* predictions, we analyzed RNA G-quadruplex sequencing (rG4-seq) data from an earlier investigation^50^, which couples rG4-meditated reverse transcriptase stalling with high-throughput sequencing to map transcriptome-wide rG4s in Hela cells treated with the rG4 stabilizer pyridostatin (PDS)^50^. PDS treatment leads to an enrichment of rG4-seq peaks around splice sites (**Figures S1H** and **S1I**), with a strong enrichment of rG4s flanking splice sites of CREs (**Figures 1I** and **1J)**.

Taken together, our analysis demonstrates that rG4 motifs are enriched around splice sites of CREs and are depleted from splice sites of CIEs. These results suggest that rG4s, that have been suggested to be more stable at lower temperatures^25,51^, could dynamically control the accessibility of splice sites, thereby serving as widespread regulators of temperature-dependent alternative splicing.

### G4 stabilizers effectively repress the inclusion of cold-repressed exons

To address a global role of rG4s in controlling temperature-dependent alternative splicing, we correlated the predicted G4 scores with the changed exon inclusion levels (delta PSI) upon cold or heat shock treatment in HEK293T or Hela cells. Interestingly, we observed that the predicted G4 score around 5’-splice sites of temperature-sensitive exons negatively correlated with the cold-induced PSI change in both Hela and HEK293T cells and found a slightly weaker positive correlations with the delta PSI upon heat shock (**Figure 2A**). As rG4s tend to be more stable at low temperatures^25,51^, this result suggests a general negative correlation between stability of rG4s around splice sites and the inclusion of the exon. To test this hypothesis, we first overlapped the identified CREs in Hela and HEK293T to obtain a high-confidence set of exons and to exclude exons with cell type-specific regulations, as splicing regulation through RNA structures could act independently of the *trans*-acting environment. In total, 1065 common CREs were identified in Hela and HEK293T cells and 380 (35.7%) of them contained predicted rG4s (score > 1) around their splice sites (**Figure 2B** and **Table S1**), pointing to a general, cell-type independent splice-regulatory function of rG4s. Next, we selected three candidate genes containing cold repressed-cassette exons with predicted rG4s around their splice sites, namely CDK4, IQSEC1 and FKBP15 (**Figures 2C**). These three genes have distinct functions in controlling cell cycling^52^, phosphoinositide metabolism^53^, and cytoskeletal organization^54^, respectively. RNA-Seq analysis unveiled temperature sensitivity of their inclusion, exhibiting elevated PSI levels at higher temperatures in both cell lines (**Figures 2D, S2A,** and **S3A**). Sequence alignments showed that the G4 motifs surrounding CREs of these genes were conserved cross multiple mammals, indicating their importance across evolution (**Figures 2E**, **S2B,** and **S3B-S3C**).

**Figure 2.**
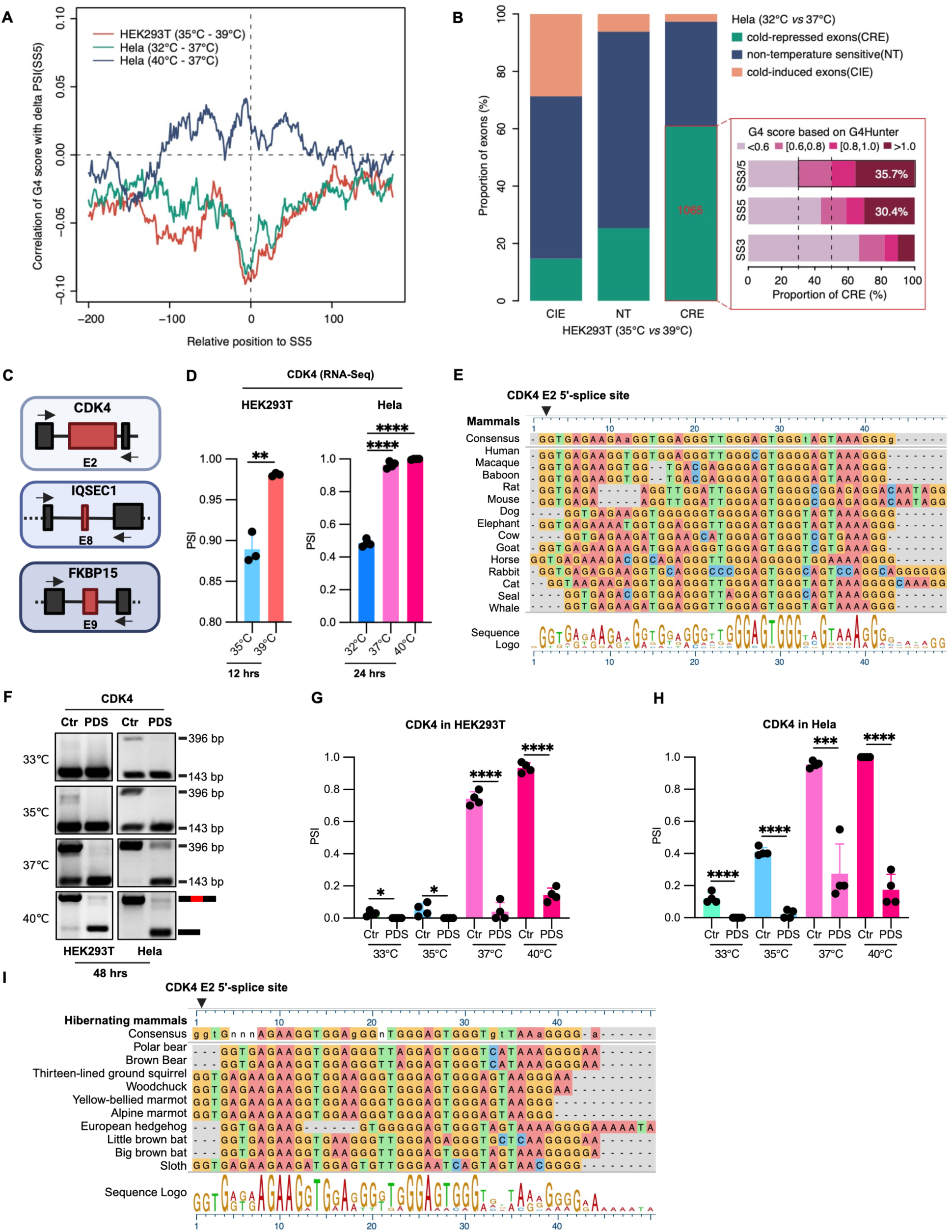
G4 stabilizers reduce the inclusion of cold-repressed exons (CREs). (A) Correlation of 5’-splice site G4 scores with delta PSI values after cold shock (HEK293T (35°C-39°C) and Hela (32°C-37°C)) or heat shock (Hela (40°C-37°C)). (B) Intersection of CIE, NT and CRE exons comparing HEK293T and Hela cells (left panel), and proportions of the shared CREs containing sequences with different G4 scores (right panel). SS3/5 shows the maximum G4 scores for either the SS3 or the SS5 for each exon. (C) Schematic depicting the position of CRE in CDK4, IQSEC1 and FKBP15. Arrows indicate RT-PCR primers. (D) PSI of CRE E2 in CDK4 determined by RNA-Seq analysis in HEK293T and HeLa cells (See table 1 and bioinformatics Method). PSI: percent spliced in. (E) Evolutionary conservation of a G-rich motif in the CRE E2 of CDK4 across multiple mammals (see bioinformatics method). (F) PSI of CRE E2 in CDK4 treated with DMSO or PDS at different temperatures in Hela and HEK293T cells. HEK293T or Hela cells were treated with 10 µM PDS and cultured at 33°C, 35°C, 37°C, and 40°C for 48 hrs (n=4), followed by RT-PCR. A representative gel image is shown. PCR products and sizes are indicated on the right. (G) and (H) Quantification results of (F), where n=4. (I) Evolutionary conservation of a G-rich motif in the CRE E2 of CDK4 across multiple hibernating mammals.

We then proceeded to validate the temperature sensitivity of splicing among these CREs and further investigated the impact of stabilizing rG4s on exon inclusion of these CREs. Upon treating HEK293T and Hela cells with the widely used quadruplex stabilizer PDS^55^, we measured the splicing levels of CREs from these three genes at different temperatures (33°C, 35°C, 37°C and 40°C) (**Figures 2F-2H**, **S2C-S2E,** and **S3D-S3F)**. Among the three candidates, CDK4 underwent the most pronounced splicing alterations following cold shock, displaying a gradual decrease in PSI from over 90% at 40°C to less than 10% at 33°C in both HEK293T and Hela cells (**Figures 2F-2H**). The inclusion of CRE E2 in CDK4 alters its N-terminal region, crucial for ATP and D-type cyclin binding, which confers its catalytic activity^56^. Within IQSEC1, the eighth exon (E8) encodes distinct segments in the C-terminal region of the protein. E8 exhibited temperature-sensitive splicing changes, with high PSI levels at 40°C dropping to less than 10% at 33°C (**Figures S2C-S2E**). While E9 of FKBP15 showed significantly decreased PSI in Hela cells following cold shock, the changes in PSI were less pronounced in HEK293T cells (**Figures S3D-S3F**), likely due to differential expression of *trans*-acting factors between cell lines. In line with our hypothesis, PDS treatment almost completely abolished the inclusion of CDK4’s E2 across the tested temperatures (**Figures 2F-2H**), and substantially reduced CRE inclusion in IQSEC1 and FKBP15 at both low and high temperatures in HEK293T and Hela cells (**Figures S2C-S2E** and **S3D-S3F**). To further examine the potential importance of the G-rich elements adjacent to these three confirmed CREs in mammals with a broader range of physiological body temperatures, we aligned the respective sequences from hibernating mammals, whose body temperatures decrease by more than 5°C during hibernation. G-rich elements were conserved across these hibernating species (**Figures 2I, S2F,** and **S3G-3H)**, implying a physiological role in regulating alternative splicing events during hibernation.

Collectively, our results underscore the robust impact of G4 stabilizers in suppressing the inclusion of CREs across different cell lines and temperatures. Furthermore, our sequence alignments show that the G-rich elements neighboring three validated CREs are conserved across mammals, including hibernating species. These findings together highlight the potential role of rG4s as modulators of temperature-dependent alternative splicing, thus acting as RNA-based temperature sensors in the mammalian body temperature range.

### Identification of G-rich elements responsible for temperature-dependent RBM3 splicing

We next aimed to further substantiate the connection between G-rich elements and temperature-sensitive alternative splicing. We focused on the cold-inducible and neuroprotective protein RBM3^57,58^, which is controlled by temperature-regulated alternative splicing of a poison exon (exon 3a) leading to heat-induced NMD^46,47^. In RNA-Seq data, RBM3’s exon 3a inclusion level remained consistently below 0.1 at or below 37°C. However, there was a significant increase observed at 39°C or 40°C in both HEK293T and Hela cells (**Figure 3A**). Furthermore, we identified putative rG4 motifs around the 3’ and 5’-splice sites of this cold-repressed poison exon, as detailed in the preceding analysis (**Figure S4A**).

**Figure 3.**
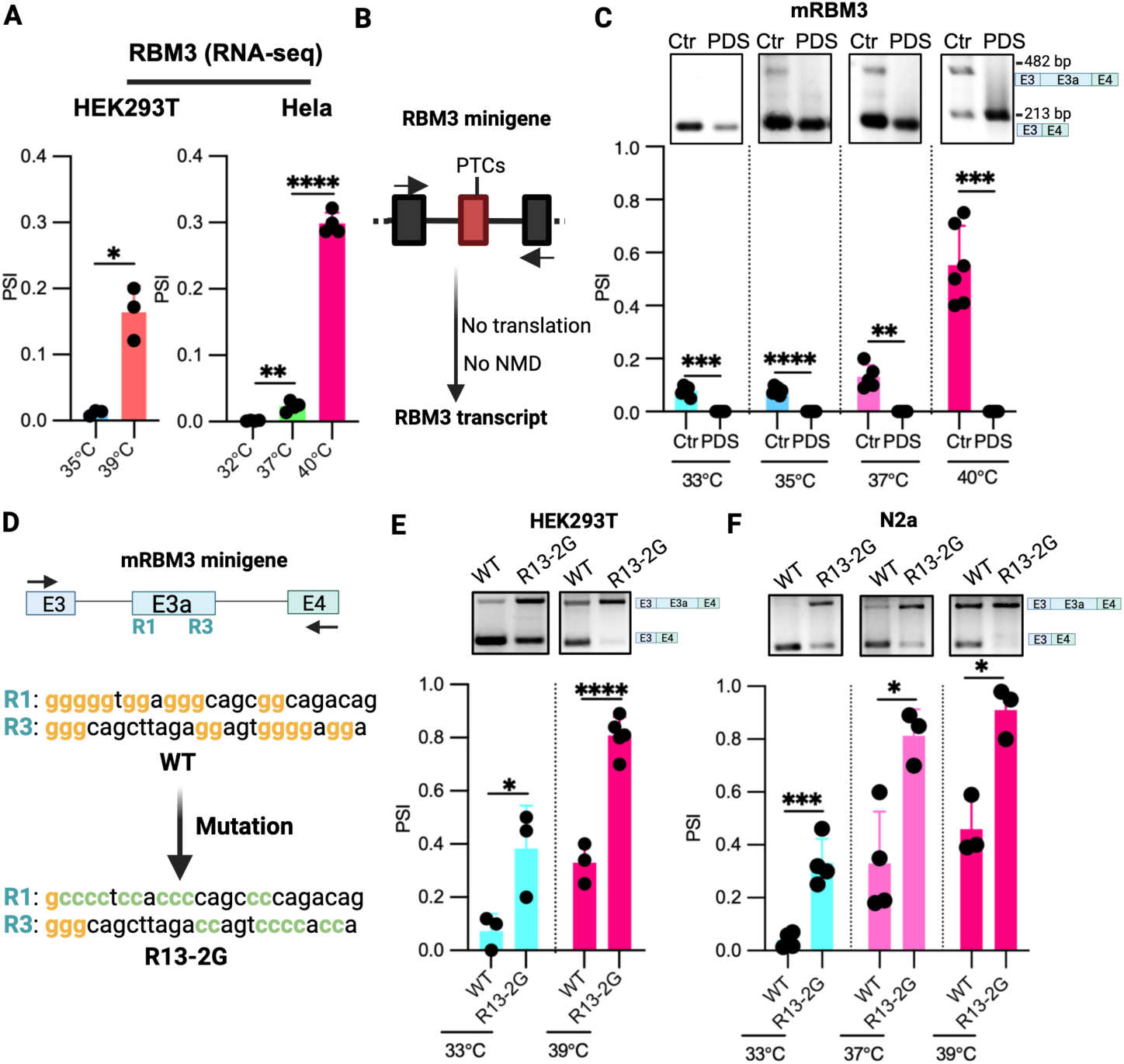
G-rich elements repress exon 3a splicing in RBM3 minigene. (A) PSI of RBM3 exon 3a in HEK293T and Hela RNA-Seq datasets from different temperatures. (see Figure 1 and bioinformatics Method). (B) Schematic of the RBM3 minigene designed to prevent translation to allow analysis of exon inclusion independent of NMD. (C) Exon 3a inclusion in the mRBM3 minigenes (mRBM3) after G4 stabilizer PDS or control treatment. Minigene-transfected HEK293T cells were incubated at the indicated temperatures for 24 hrs. Upper gels depict representative minigene specific RT-PCR results, and the lower part shows quantified results (n=6 for 35°C and 40°C, n=5 for 33°C and 37°C). (D) Schematic illustrating the position of the G-rich elements R1 and R3 in the mRBM3 minigene and the sequence of the R13-2G mutant. (E) and (F) Splicing level of RBM3 exon 3a in WT and R13-2G mutant at different temperatures in both HEK293T (E) and N2a cells (F). Upper gels depict representative RT-PCR results, and the lower portion shows quantified results (n ≥3).

Since RBM3 transcripts containing exon 3a are extremely efficiently degraded via the NMD pathway^46,47^, we cloned human RBM3 (hRBM3) and mouse RBM3 (mRBM3) minigenes consisting of exon 3, exon 3a, exon 4, and the introns flanking exon 3a. The RNA produced by the two minigenes is not translated, and therefore is not degraded by NMD (**Figure 3B**). In both hRBM3 and mRBM3 minigenes, the splicing of exon 3a maintained temperature sensitivity, and PDS treatment almost fully blocked exon 3a inclusion at various temperatures (**Figures S4B** and **3C**).

To explore the impact of G-rich elements near splice sites of RBM3 exon 3a, we introduced mutations to each G-rich element in five regions: two around the splice sites (R1 and R3), one within exon 3a (R2) and two in the downstream introns (R4 and R5) (**Figures S4A** and **S4C**). Interestingly, mutations or deletions of G-rich elements in R1 and R3 increased exon 3a inclusion in HEK293T (**Figure S4D)** and mouse N2a cells (**Figure S4E)**. Using MaxEntScan^59^, we further excluded the possibility that the effect of the mutations in promoting exon 3a inclusion was caused by increasing 3’-splice site strength (**Figure S4F)**.

Collectively, the mutagenesis assay supports the hypothesis that G-rich elements in regions R1 and R3 contribute to RBM3 exon 3a alternative splicing through masking the closely adjacent splice sites. We also found that the four stretches of consecutive G residues in both R1 and R3, that potentially form an rG4 structure, are evolutionarily conserved in mammals, indicating their importance in RBM3 gene regulation (**Figures S4G** and **S4H)**. Furthermore, we observed the conservation of R1 and R3 among hibernating animals, suggesting the potential significance of these two elements in regulating RBM3 expression during hibernation (**Figures S4I** and **S4J)**. As we hypothesized that G-rich elements in R1 and R3 could individually mask one of the splice sites, their combined mutation (R13-2G) might result in even stronger promotion on RBM3 exon 3a inclusion (**Figure 3D)**. Indeed, R13-2G increased isoforms containing exon 3a in both HEK293T cells and N2a cells (**Figures 3E** and **3F**) with a stronger effect than each single mutant **(Figure S4E)**, again confirming that these G-rich sequences repress exon 3a inclusion.

### Biophysical assays reveal RNA G-quadruplex formation in RBM3

As our results demonstrate that G-rich elements around RBM3 exon 3a governed inclusion of this exon, likely through modulating splice site accessibility, we aimed to provide direct evidence that these sequences represent rG4 structures. To this end, we used an RNA oligonucleotide corresponding to the sequence in R1 and a mutant version (R1-mutant), in which the essential G nucleotides are replaced with Cs (**Figure 4A**). We then measured rG4 structure formation with circular dichroism (CD) spectroscopy, one of the standard methods to detect rG4s (**Figure 4B**)^60^. For the R1 RNA, we observed a spectrum showing a clear and strong positive signal peak at 265 nm and a negative signal peak at 240 nm, which is the specific CD spectrum for a parallel rG4^60^. This signal was dependent on KCl^60^ and the signal was not observed for the R1-mutant, supporting the notion that the WT sequence formed an rG4 structure *in vitro* (**Figure 4C**). Furthermore, we found that the intensity of the peak was temperature-sensitive at low potassium concentration (0.1 mM) (**Figures 4D** and **4F**), while it showed less temperature-sensitivity at high potassium concentration (50 mM) (**Figures 4E** and **4F**). Interestingly, at low potassium concentration (0.1 mM), the temperature-controlled changes in signal intensity were reversible when the RNA was first incubated at 40°C and then cooled down to 33°C (**Figures 4D** and **4F**), showing a dynamic reaction to changes in the physiological temperature range. It should be noted that the potassium concentrations used *in vitro* cannot be directly compared with the conditions in cells, as additional factors contribute to controlling the stability of rG4 structures in the cellular environment.

**Figure 4.**
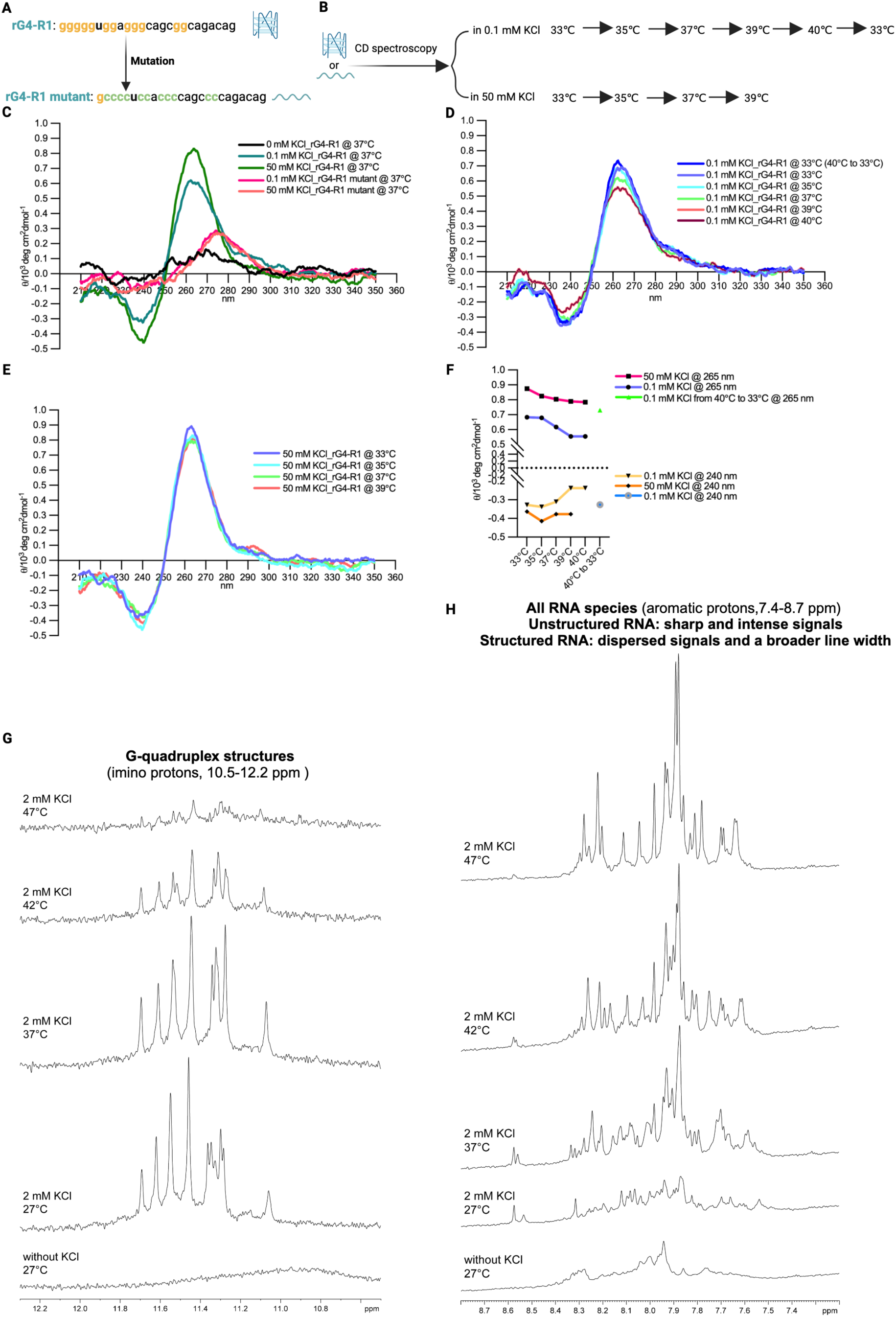
Biophysical assays demonstrate temperature and potassium dependent G-quadruplex formation of RBM3 R1 *in vitro*. (A) Sequences of synthesized WT and mutant rG4-R1 RNA oligos. (B) A schematic of biophysical CD spectroscopy assays. (C) Circular dichroism (CD) spectrum signal of rG4-R1 and its mutant in different potassium concentrations at 37°C. X-axis: the wavelength of light; y-axis: the magnitude of the CD signal. Signal peak at 265 nm and 240 nm: parallel rG4 structure. (D) CD signal of rG4-R1 at different temperatures in low KCl concentration. (E) CD signal of rG1-R1 at different temperatures under high KCl concentration. (F) A summary of CD signal peaks at 265 nm and 240 nm of rG4-R1 at various temperatures under both high and low concentrations of KCl. (G) and (H) 1D ^1^H NMR spectrum showing the imino region (G, peaks indicating structured rG4s) and aromatic protons (H, as a measure for unstructured RNA, see text for details) confirming that the rG4-R1 sequence forms a temperature sensitive rG4 *in vitro*.

Finally, to further validate the presence of an rG4 structure, we recorded a 1D ^1^H NMR spectrum of the rG4-R1 RNA at different temperatures and potassium concentrations (**Figures 4G** and **4H**). We first focused on the imino region (10-16 ppm), in which exchange-protected imino protons due to hydrogen-bonding are clear indicators of RNA secondary structure. Imino protons involved in Watson-Crick base-pairing are found in a region of 12-15 ppm, whereas imino protons of G-quadruplex structures show signals in a very narrow region of 10.5-12.2 ppm^61^. In the absence of potassium, no imino signals were observed indicating the lack of a stable RNA structure. Upon addition of potassium, however, a set of well-dispersed imino signals clearly indicated the presence of a quadruplex structure at 27°C. These signals decreased at elevated temperatures, especially between 37°C and 42°C, indicating a partial loss of the quadruplex structure in this temperature range, and were almost completely lost at 47°C (**Figure 4G)**. Furthermore, the NMR signal of aromatic protons that show signals for all RNA species including unstructured and structured RNA gives a more complete picture (**Figure 4H**). Unstructured RNA results in sharp and intense signals whereas structured RNA contributes more dispersed signals with a broader line width due to the slower molecular tumbling of the larger structure. This region of the ^1^H spectrum showed the absence of unstructured RNA at 27°C and a strong increase of unstructured RNA already at 37°C, which was further increased at 42°C and 47°C (**Figure 4H)**. These results together strongly suggest the formation of an rG4 structure, which dynamically reacts to temperature changes in the mammalian body temperature range, thus representing a mammalian RNA thermometer.

### Mutations of the G-rich element abolish the effect of G4 stabilizers on mRBM3 exon 3a exclusion

After demonstrating that the G-rich sequence around splice sites of RBM3 exon 3a can form rG4 structures *in vitro*, we aimed to confirm their relevance for RBM3 exon 3a splicing in cells. Thus, we turned to K^+^ as an endogenous G4 stabilizer, and tested the impact of KCl on the mRBM3 minigene and the rG4 double mutant (R13-2G) in HEK293T and N2a cells. In the WT mRBM3 minigene, KCl treatment significantly decreased RBM3 exon 3a inclusion at high temperatures, while it had no effect at low temperatures (**Figures 5A-5D)**. This suggests that higher stability of the rG4 structures at lower temperatures leads to reduced dependence on potassium. In contrast, exon 3a inclusion in the double mutant (R13-2G) did not significantly change upon KCl treatment at both low and high temperatures (**Figures 5A-5D)**. To further investigate the impact of stabilizing rG4s on exon 3a splicing at normal physiological temperature (37°C) in a neuronal context, we treated HT22 cells, a mouse hippocampal neuronal cell line, with KCl. Consistently, KCl promoted exon 3a skipping in the WT mRBM3 minigene, while it had no effect on the R13-2G mutant minigene (**Figures 5E** and **5F**). Furthermore, to mimic the pathological brain injury condition, we treated glutamate-stimulated HT22 cells with the broad potassium channel blockers 4-AP and AFP (amifampridine) to pharmacologically increase intracellular potassium levels and examined exon 3a splicing at 37°C. Both 4-AP and AFP significantly reduced exon 3a inclusion in the WT, but not in the R13-2G minigene (**Figures 5G** and **5H**). These results together strongly suggest that rG4s in RBM3 exon 3a are KCl-responsive and temperature-dependent RNA elements, that, by masking splice sites, control exon inclusion in cells.

**Figure 5.**
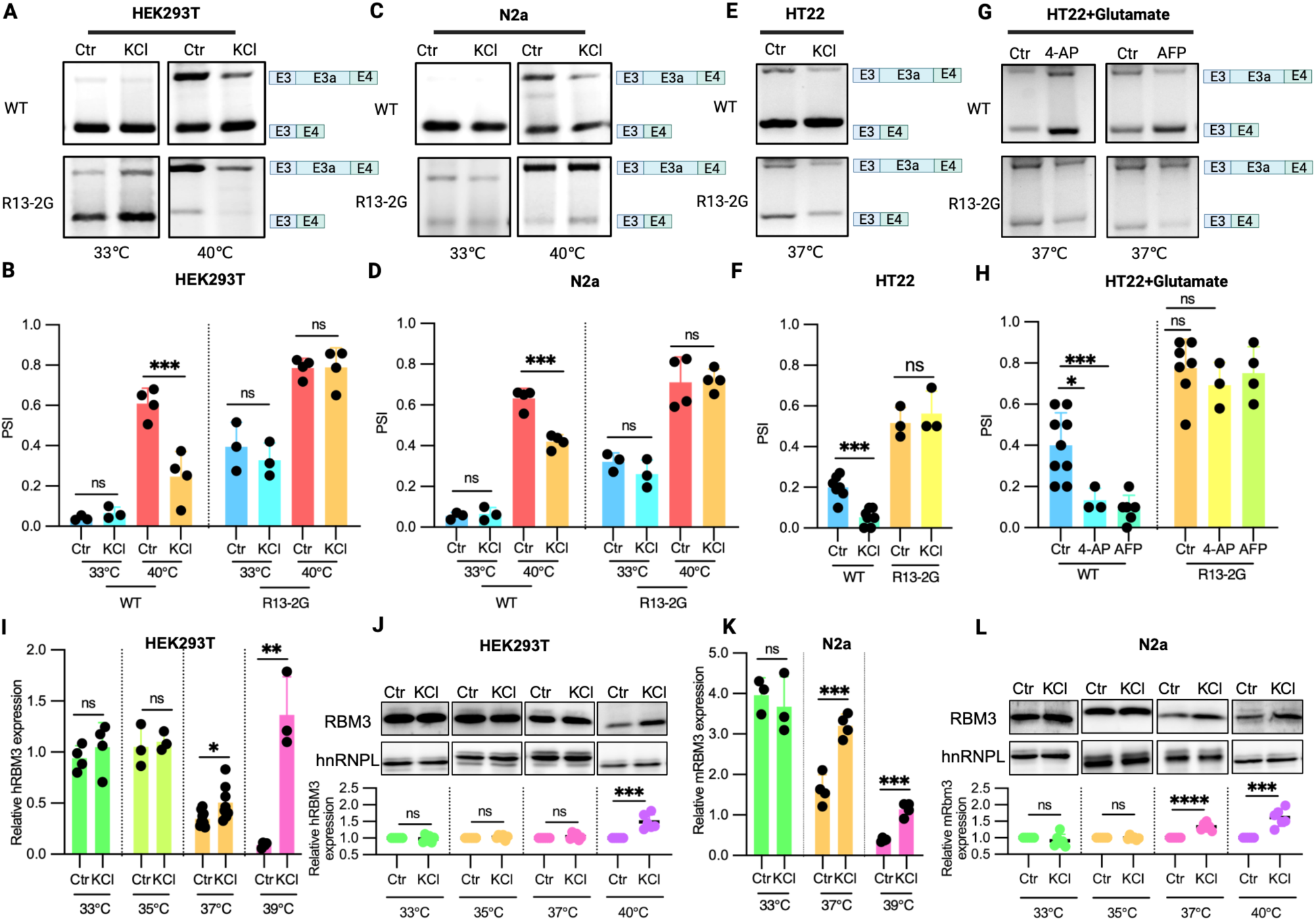
RBM3 rG4 elements control RBM3 levels in response to G4 stabilizers. (A-D) Splicing level of RBM3 exon 3a in the WT and R13-2G double mutant mRBM3 minigene after KCl and control treatment at 33°C and 40°C in both HEK293T (A and B) and N2a (C and D) cells (see Method, n ≥*3*). (E) and (F) KCl treatment as in (A-D) in HT22 cells at 37°C (see Method, n ≥*3*). (G) and (H) Splicing level of RBM3 exon 3a after 4-AP and AFP treatment in glutamate-stimulated HT22 cells at 37°C (see Method, n ≥3). AFP: amifampridine. 10 μM of 4-AP and AFP were used. (I-L) RBM3 expression after KCl and control treatment at 33°C, 35°C, 37°C or 39°C, observed for both mRNA and protein levels in HEK293T (I and J) and N2a (K and L) cells. The left panel results are derived from qPCR, and the right panels are WB results. Below the gels, quantifications using HNRNPL as loading control are shown (see Method, n ≥3).

### Increasing endogenous potassium promotes endogenous RBM3 expression

We then used KCl to stabilize rG4s in RBM3 exon 3a and investigated the impact on endogenous RBM3 expression. In line with our hypothesis, KCl treatment increased RBM3 mRNA and protein expression at high (≥37℃) but not at low (<37℃) temperatures in HEK293T and N2a cells (**Figures 5I-5L**). Collectively, these findings indicate that both low temperatures and potassium ions are capable of stabilizing rG4 structures, thereby increasing endogenous RBM3 expression.

### 4-AP protects neurons from hemin-induced damage in an RBM3-dependent manner

It has been well established that RBM3 safeguards neurons and ameliorates phenotypes in prion disease and hypoxic-ischemic brain injury mouse models by facilitating neuronal structural plasticity, preventing cell death and promoting neurogenesis^57,58^. Here, we proceeded to test whether increasing endogenous RBM3 expression could protect from neuronal damage in a hemin-induced hemorrhagic stroke model^62^. In this model, hemin, a derivative of heme – an essential component of hemoglobin in red blood cells – mimics the oxidative stress and neuroinflammation of hemorrhagic stroke. We first used a cell culture model of hemin-induced cell death and observed that hemin treatment led to a notable reduction in intracellular potassium concentration (**Figure 6A**) and prompted exon 3a inclusion, as evidenced by a higher ratio of exon 3a to RBM3 mRNA (**Figures 6B** and **6C**), ultimately resulting in an almost two-fold reduction of RBM3 expression (**Figure 6D**). We then pharmacologically increased intracellular potassium levels via the potassium channel blocker 4-AP. 4-AP treatment significantly elevated intracellular potassium levels in a dose-dependent manner (**Figure 6A**) and concurrently led to a gradual reduction in the exon 3a to total RBM3 ratio (**Figures 6B** and **6C**) and an increase in RBM3 mRNA expression (**Figures 6D**). Consequently, RBM3 protein expression was substantially increased following 4-AP treatment (**Figures 6E-6G**), reaching a nearly fivefold protein elevation compared to hemin treatment alone (**Figures 6E-G**). Phenotypically, the hemin-induced cell death rate decreased (**Figures 6H** and **6I**), from 25% to 5%, and cell viability increased significantly and substantially following 4-AP treatment (**Figure 6J**). Knocking down RBM3 (**Figure 6K**) with siRNA post-4-AP treatment abolished these phenotypic effects (**Figures 6L-6N**), underscoring the pivotal role of RBM3 in mediating the neuroprotection conferred by 4-AP. To further validate RBM3’s protective role against hemin-induced neuronal damage, we overexpressed RBM3 in HT22 cells (**Figures 6O** and **6P**). This led to an increase of cell viability from 60% to 80% (**Figure 6Q**) and a notable decline in cell death from 16% to 7% (**Figures 6R** and **6S**) upon hemin treatment.

**Figure 6.**
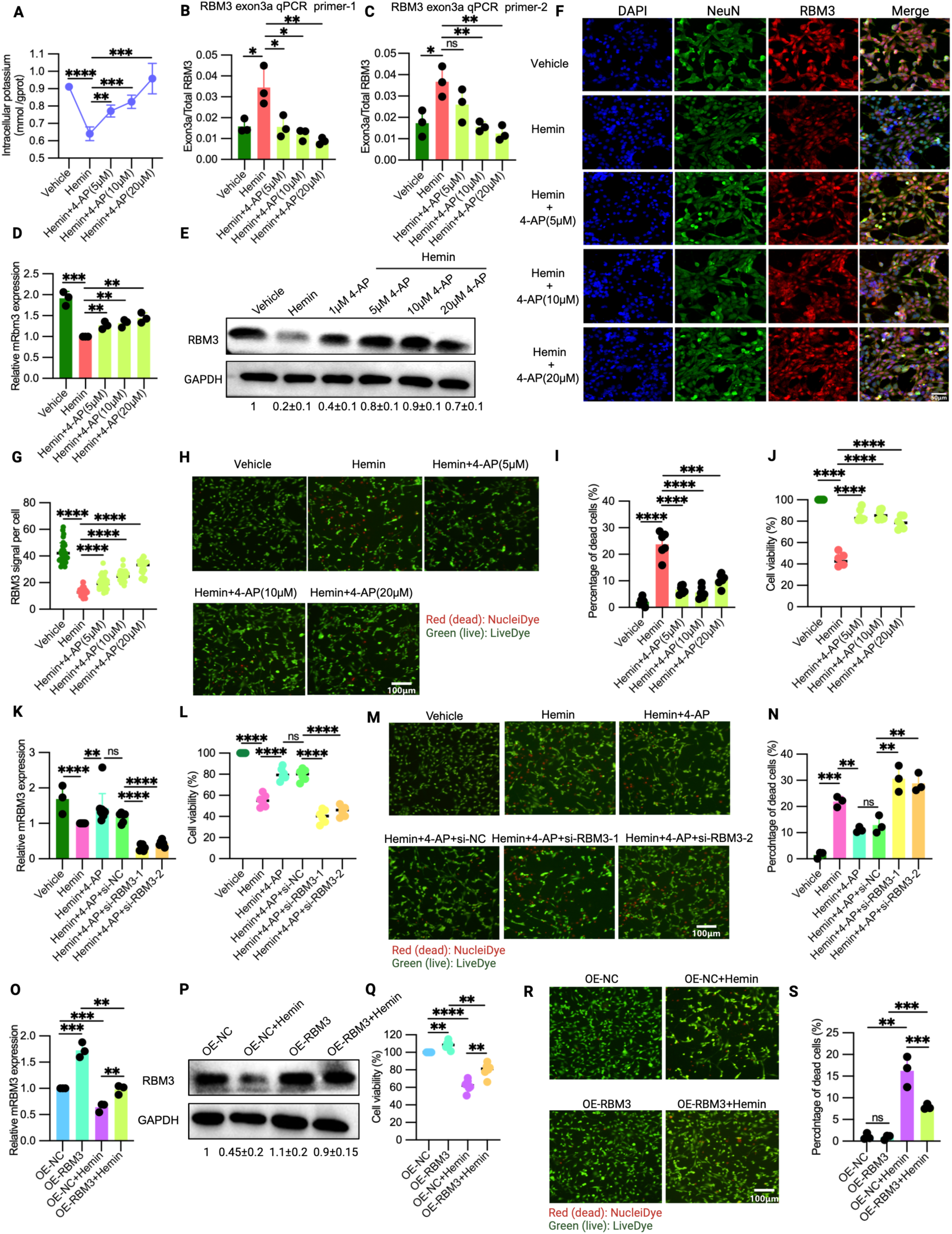
4-AP confers RBM3-mediated protection against neuronal damage in a hemin-induced hemorrhage cell model. (A) Intracellular potassium levels after 4-AP and control treatment in hemin-exposed HT22 cells (see Method, n=4,). (B) and (C) Levels of RBM3 exon 3a inclusion after 4-AP and control treatment in HT22 cells treated as in (A). Two pairs of qPCR primer (RBM3 exon3a qPCR primer-1(B) and RBM3 exon3a qPCR primer-2 (C)) targeting the exonic junction region of exon 3a and its upstream and downstream exons, and the exonic regions directly adjacent to the 3’ and 5’-splice sites of exon 3a, respectively) were used to quantify exon 3a inclusion (n=3), respectively. (D) Endogenous RBM3 mRNA expression after treatments as in (A) (n=3). (E) Endogenous RBM3 protein expression as treatments in (A), revealed by Western blot analysis. Below the gel, quantification (mean ± SD) using GAPDH as loading control is shown (n=3). (F) Immunostaining of DAPI, NeuN, RBM3 and merged images as treatments in (A) (n=3). (G) RBM3 expression was quantified from (F) (n=3). Immunostaining data is depicted in (F), with quantification in (G) (n=3). (H) and (I) Cell death in HT22 cells as treatments in (A). Images stained for dead/live cells are shown in (H), with quantification in (I) (see Method, n=3). (J) Cell viability as treatments in (A), as shown by CCK-8 assays (see Method, n=6). (K) Relative RBM3 mRNA expression post-siRNA upon 4-AP treatment in hemin-exposed HT22 cells. Two independent si-RNAs against RBM3 were used (see Method, n≥3). (L) Cell viability as treatments in (K), shown by CCK-8 assay (n=6). (M) and (N) Percentage of dead cells as treatments in (K). Stainings for dead/live cells are provided in (M), with quantification in (N) (see Method, n=3). (O) and (P) Relative RBM3 expression after lentiviral overexpression in hemin- and control-treated HT22 cells. RBM3 mRNA expression was assessed by qPCR in (N) (n=3), while RBM3 protein expression was measured by Western blot in (O). Below the gel, quantification relative to GAPDH is shown (see Method, n=3). (Q) Cell viability as treatments in (O), shown by CCK-8 assay in HT22 cells (n=6). (R) and (S) The percentage of dead cells as treatments in (O). Stained images for dead/live cells are presented in (R), with quantification in (S) (see Method, n=3).

### 4-AP protects neurons from neuronal damage in a subarachnoid hemorrhage mouse model

To further address 4-AP’s efficacy and potential for clinical use, we subsequently evaluated the impact of 4-AP *in vivo* using a subarachnoid hemorrhage (SAH)^63,64^ mouse model (**Figure 7A)**. Given that the most impacted area in individuals with SAH is in close proximity to the bleeding site, we directed our attention to the cerebral cortex surrounding the affected region (**Figure 7B)**. Intraperitoneal administration of 4-AP resulted in a significantly elevated intracellular potassium level in the cerebral cortex (**Figure 7C**), upregulation of endogenous RBM3 mRNA expression (**Figure 7D)** and decreased RBM3 exon 3a inclusion (**Figures S5A, 7E,** and **S5B**). RBM3 protein expression was also augmented, as demonstrated by immunostaining (**Figures 7F** and **S5C**). Notably, the increased RBM3 expression following 4-AP treatment correlated with decreased neuronal apoptosis (**Figure 7G**) from ∼30% to 15%, as indicated by TUNEL staining^65^ (**Figure I)**, and increased neuronal count (**Figure 7H**) shown by neuronal nuclear protein (NeuN) staining^66^ (**Figure 7I)**. Hematoxylin and eosin (HE) staining also revealed that 4-AP treatment significantly ameliorated the severity of cortical spongiosis^67^ resulting from SAH (**Figures 7J** and **7K**). 4-AP administration also prevented neuronal damage, which is apparent as eosinophilic necrotic neurons with aberrant morphology characterized by cell body shrinkage, darkly stained pyknotic nuclei, and intensely stained eosin, with a significant increase in neuronal No. from around 20 to 60 (**Figures 7J** and **7L**). Further, Nissl staining^68^ demonstrated increased cell counts with normal morphology following 4-AP treatment compared to the SAH control group (**Figures S5D** and **S5E**). In addition, a significant increase in the intensity of Nissl substance was observed post 4-AP treatment, effectively counteracting the loss of Nissl substance induced by SAH (**Figures S5D** and **S5F**). The higher staining intensity of Nissl substance following 4-AP treatment may suggest increased metabolic activity and possibly increased protein synthesis. Finally, 4-AP treatment markedly improved mouse behavioral performance, manifested by increased modified Garcia scores^68,69^ (**Figure 7M**) and enhanced latency to fall in the Rotarod test^70–72^ (**Figures 7N** and **7O**).

**Figure 7.**
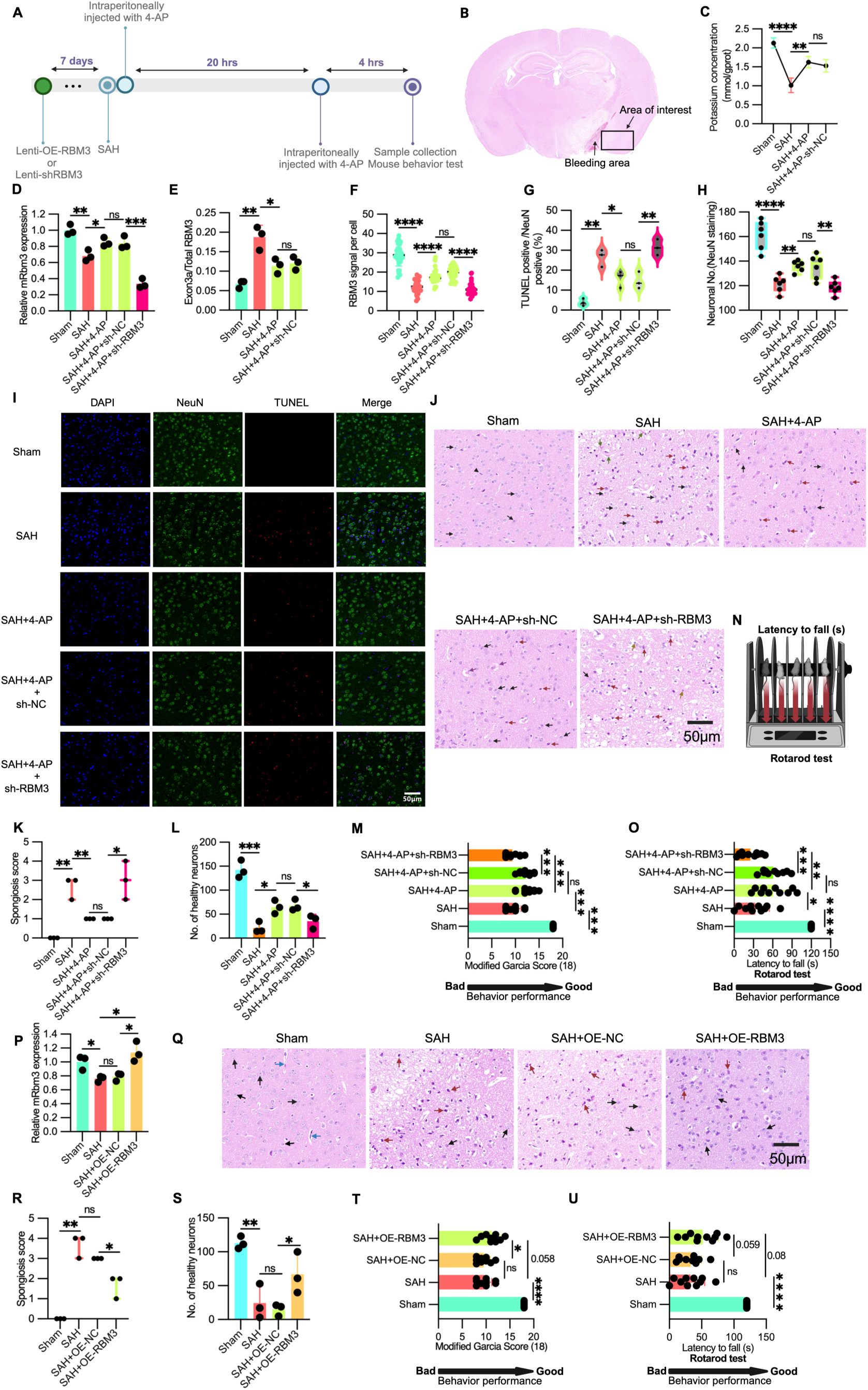
4-AP mitigates neuronal damage in a mouse model of subarachnoid hemorrhage (SAH).

(A) Timeline of *in vivo* mouse experiments.

(B) Mouse cerebral cortex for the following investigation (D-L) and (P-S).

(C) Intracellular potassium levels in cortical brains of the SAH and sham mouse model *in vivo* after 4-AP and control administration (see Method, n=3 mice).

(D) RBM3 mRNA expression of the SAH injected with lenti-shRBM3 and lenti-NC and sham mouse model *in vivo* after 4-AP and control administration, shown by qPCR (see Figure 7A and Method, n=3 mice).

(E) Inclusion level of RBM3 exon 3a in (B) of the SAH and sham mouse model *in vivo* after 4-AP and control administration. RBM3 Exon 3a expression was quantified with RBM3 exon 3a qPCR primer-1 normalized with mGAPDH (n=3 mice).

(F) RBM3 protein expression of the mice treated as (D), shown by RBM3 signal per cell in RBM3 immunostaining (n=3 mice).

(G) Apoptotic cells of the mice treated as (D). The data was quantified from (I) (n=3 mice).

(H) Neuronal count of the mice treated as (D). Data was quantified from NeuN signal-positive cells in (I) and Figure S5C (n=6 mice).

(I) Representative immunostaining images of DAPI, NeuN and TUNEL in (B) of the mice treated as (D).

(J) Representative images of HE staining in (B) of the mice treated as (D). Representative healthy neurons are denoted by black arrows, damaged neurons by red arrows, red blood cells in the capillary by green arrows, healthy glia by arrowheads, damaged glia by yellow arrows, and macrophages by purple arrows (n=3 mice).

(K) Spongiosis score in (B) of the mice treated as (D), quantified from hematoxylin and eosin (HE) staining (J) (n=3 mice).

(L) The No. of healthy neurons in (B) of the mice treated as (D), quantified from hematoxylin and eosin (HE) staining (J) (n=3 mice).

(M) Modified Garcia score of the mice treated as (D) (see Method^68^, n=10 mice).

(N) Schematic representation of the Rotarod Test (see Method).

(O) Latency to fall of the mice treated as (D) in the Rotarod Test (see Method, n=10 mice).

(P) RBM3 mRNA expression of the SAH injected with lenti-OE-RBM3 and lenti-NC and sham mouse model *in vivo* (also see Figure 7A and Method, n=3 mice).

(Q) Representative HE staining images in (B) of the mice treated as (P). Representative healthy neurons are denoted by black arrows, damaged neurons by red arrows and endothelial cells in the capillary by blue arrows (n=3 mice).

(R) Spongiosis score in (B) of the mice treated as (P), shown by HE staining in (Q) (n=3 mice).

(S) The No. of healthy neurons in (B) of the mice treated as (P), indicated by HE staining in (Q) (n=3 mice).

(T) Modified Garcia Score of the mice treated as (P) (see Method^68^, n=10 mice).

(U) Latency to fall of the mice treated as (P) in the Rotarod Test (see Method, n=10 mice).

To confirm whether the beneficial effects of 4-AP treatment in the SAH mouse model are dependent on increased RBM3 expression, we utilized an shRNA lentivirus to knock down RBM3 *in vivo* prior to SAH and 4-AP treatment (**Figures 7A, 7D, 7F,** and **S5C**). Consequently, this resulted in increased neuronal apoptosis^65^ (**Figures 7G** and **7I**), reduced neuronal count (**Figure 7H**), as evidenced by NeuN^66^ staining (**Figure 7I)**, elevated spongiosis score (**Figures 7J** and **7K**) and decreased healthy neurons (**Figures 7J** and **7L**) shown by HE staining (**Figures 7J)**, decreased cells with normal morphology indicated by Nissl staining^68^ (**Figures S5D** and **S5E**), and diminished intensity of Nissl substance (**Figures S5D** and S**5F**) compared to 4-AP treatment alone. Moreover, the improved mouse behavior elicited by 4-AP treatment was nullified by RBM3 knockdown, as evidenced by decreased modified Garcia scores^68,69^ (**Figure M**) and reduced latency to fall in the Rotarod test^70–72^ (**Figures 7N** and **7O**).

To further illustrate the neuroprotective efficacy of RBM3 against SAH, we augmented RBM3 expression *in vivo* in mice via lentivirus-mediated RBM3 overexpression prior to SAH (**Figures 7A** and **7P**). Heightened RBM3 levels were markedly correlated with diminished spongiosis severity (**Figures 7Q** and **7R**), increased healthy neurons (**Figures 7Q** and **7S**), increased number of cells with normal morphology (**Figures S5G** and **S5H**), and increased intensity of Nissl substance^68^ (**Figures S5G** and **S5I**). RBM3 upregulation also significantly elevated Garcia scores^68,69^ in mice with SAH (**Figure 7T**) and modestly improved performance in the Rotarod test^70–72^, as evidenced by increased latency to fall (**Figure 7U**).

Taken together, our results show that G-rich sequences around splice sites of RBM3 exon 3a form stable rG4 structures at low temperatures (33℃ and 35℃) or high potassium concentration that repress exon inclusion, resulting in increased expression of the neuroprotective RBM3 by preventing NMD. This regulation may be exploited in clinical settings, as we show that potassium-mediated increase in RBM3 acts neuroprotective *in vivo*. Since we found that a substantial percentage of CREs contain rG4 motifs, rG4s may serve as widespread physiological thermo- and potassium-sensors regulating alternative splicing in mammalian cells, which may be targeted in new therapeutic approaches in diverse conditions.

## Discussion

Alternative splicing is an essential gene regulatory mechanism shaping the complex transcriptome in various conditions across different mammalian cells. Temperature was shown to alter splice site choice through changing the activity of *trans*-acting factors, whose expression may vary across cell types, which contributes to splicing heterogeneity. Here, we describe a mechanism that depends on the RNA itself, and could thus function in *cis*. We show that altered temperature-controlled formation of rG4 structures in the vicinity of splice sites of cassette exons controls alternative splicing. Our data show that around 10-20% (putative rG4 motifs) or even 35.7% (G4Hunter) of exons whose inclusion is repressed upon cold shock, contain potential rG4s in sequences around their splice sites, suggesting a widespread effect of rG4s as physiological thermo-sensor in mammalian cells. We suggest that temperature-(and potassium) controlled formation of rG4s alters the accessibility of splice sites thereby leading to temperature-controlled alternative splicing. Based on our *in vitro* data showing modulation of the rG4 structure in the physiological temperature range, and corresponding mutational analysis in cell culture experiments, we classify these sequences are mammalian RNA thermometers. Temperature-controlled behavior of rG4s will be additionally controlled by RBPs in the cellular environment, which may alter the exact temperature response of the RNA. However, as the temperature range with dynamic response of the RBM3 rG4 is very similar *in vitro* (CD and NMR) and in cells (exon skipping and RBM3 expression), we suggest, that at least in this case, the RNA is sufficient to act as temperature sensor and is not substantially influenced by cellular RBPs. Besides controlling accessibility to the splice sites, RNA conformational dynamics may alter the binding of additional proteins in these regions, since many RNA binding proteins (RBPs) prefer binding to single-stranded sequences, whereas others bind to double-stranded or structured RNA. In addition to directly altering splice site accessibility, we, therefore, speculate that RBPs^43,73,74^ such as hnRNPH1^47^ might preferentially bind either stable rG4s or single-stranded G-rich sequences to contribute to temperature-dependent alternative splicing of these exons.

We observed that the repressive effects of G4 stabilizer on exon inclusion are distinct across different rG4 elements. For instance, PDS can almost fully block the inclusion of CREs from CDK4 (**Figures 2F-2H)** and RBM3 minigenes (**Figures 3C** and **S4B**), but the inclusion of CREs from IQSEC1 (**Figures S2C-S2E)** and FKBP15 (**Figures S3D-S3F**) is only partially impeded by the same concentration of PDS. It has been reported that PDS has different effects on divergent rG4s^75^. One possibility is that rG4s are not naked and may associate with other *trans*-acting factors to repress inclusion of CREs. If these *trans*-acting factors are distinct for different CREs, they may respond differentially to G4 stabilizers. Another possibility is that the binding affinity of PDS to different rG4s varies due to structural differences. Quadruplex structures with distinct backbone orientations (parallel, antiparallel), different loops or additional stabilizing elements may exhibit different affinities to stabilizing ligands. These structural disparities could explain the varying responses of PDS to specific rG4s, and may also offer the opportunity to design small molecules that preferentially or specifically stabilize individual rG4s. Such specific rG4 stabilizers would provide very promising therapeutic approaches, as they could for example be used to specifically increase RBM3 expression, without affecting other rG4-dependent gene expression events. In another project, we have developed ASOs that specifically control RBM3 exon 3a inclusion thereby regulating RBM3 expression^46^. These ASOs have the clear benefit of being highly specific, but at the same time, delivering ASOs to the central nervous system requires intrathecal injection and the production of ASOs in sufficient quality and quantity for use in humans is associated with high costs. Therefore, orally available specific rG4 stabilizers could represent a promising alternative. Interestingly, our ASOs target an exonic splicing enhancer independent of the rG4s described here, suggesting that there are at least two independent mechanisms controlling RBM3 exon 3a inclusion and RBM3 expression. Combining both approaches may further increase the potential of RBM3 modulating therapeutic approaches.

KCl depolarization is known to alter alternative splicing^76^, yet mechanistic details are not fully understood. Our findings imply that at least a subset of K^+^ responsive exons are regulated by stabilizing rG4s, which is in line with the essential role of K^+^ in the formation of stable rG4 structures. This likely translates to *in vivo* situations, as maintaining potassium homeostasis is vital for cellular biochemistry^77^ and intracellular potassium imbalances, observed for example in kidney disorders^77^, vascular disease^78^ and brain disorders^79,80^ including Alzheimer’s disease^81^, may contribute to dysregulated gene expression and/or the disease phenotype. Here, we provide a molecular potential: increased intracellular potassium leads to elevated expression of RBM3, which may act neuroprotective in diverse conditions. It is interesting to note that the neuroprotective role of 4-AP in our model systems seems to be mediated primarily through increased RBM3 expression, as the beneficial effect is strongly reduced or absent upon RBM3 knockdown. This finding is somewhat surprising, given that 4-AP treatment and the ensuing increase in intracellular potassium concentration are expected to have a broader impact on gene expression. However, additional molecular targets that may contribute to the beneficial effect of 4-AP remain to be discovered. Altogether, our results lay the groundwork for promising new therapeutic avenues for neurodegenerative diseases that could benefit from elevated RBM3 expression, and more broadly, for conditions that can be treated with hypothermia.

### Limitations of our study and future work

Our study demonstrates the presence of thermos-sensing rG4 structures in sequences surrounding splice sites of cold-repressed exons and unveils a neuroprotective role of regulating temperature-dependent RBM3 splicing in cell lines and animal models. However, to precisely follow this rG4 in a temperature-dependent manner in the cellular environment in the presence of diverse RBPs that can stabilize or destabilize RNA structures, *in vivo* SHAPE, rG4-seq or in-cell NMR would be interesting for further experiments. In addition, *in vivo* RNA G4-seq could provide insights into global rG4 structures under varying temperatures. Such experiments could also reveal sequence features that render an rG4 temperature-sensitive *vs* temperature-insensitive (i.e. stable) rG4 structures. This will be very interesting to explore, as rG4s have been described as rather stable RNA structures and may therefore not have been considered potential RNA thermometers so far. Finally, mammalian body temperature is not uniform across different organs and tissues, particularly skin and testis are maintained below core body temperature levels. It will be fascinating to explore to which extent dynamic rG4s contribute to temperature-dependent gene expression patterns, for example in different tissues but also in other conditions that alter (core) body temperature, such as hibernation, exercise, fever, the circadian rhythm or ageing.

## Supporting information

Supplementary figures and figure legens

## Materials and Methods

### Cell culture and treatment

HEK293T and HT22 cells were cultured on uncoated plastic flasks or plates in DMEM medium (4.5 g glucose/L, supplied with GlutaMAX L-glutamine, Gibco,10566) supplemented with 10% fetal bovine serum (Biochrom) and penicillin-streptomycin (1:00 from a stock of 10000 U/mL, Gibco, 15140122) in 5% CO2 at 37℃ except where stated otherwise. N2a cells were cultured also on plastic flasks or plates in the medium with DMEM/opti-MEM (1:1) supplemented with 10% fetal bovine serum (Biochrom) and penicillin-streptomycin (1:00 from a stock of 10000 U/mL, Gibco, 15140122). At roughly 80% confluency, cells were sub-cultured using a 0.25% trypsin solution (1:10 dilution of 2.5% stock, Gibco,15090046) in phosphate-buffered saline. HEK293T and Hela cells were treated with 10 μM PDS at different temperatures for 48 hrs. The HEK293T or N2a or HT22 were treated with KCl (50 mM) at different temperatures for 24 hrs (for qPCR) or 48hrs (for WB). HT22 cells were treated with different dosages of 4-AP or amifampridine, and hemin (150 μM) if mentioned together for 24 hrs. For the RBM3 overexpression assay, HT22 cells were seeded into 12-well plates until the cell confluence reached 20-30%. Then, the medium of each well was replaced with complete culture medium (200 μl) containing 10 μl of virus (at a concentration of 1×10^8 TU/ml) and 8 μl of HitransG P solution (REVG005, Gekai). After 24 hours of culture, the medium was replaced to fresh complete culture medium again. Cell viability and EGFP fluorescence expression were observed under a microscope 72 hours after infection. Subsequently, the cells were cultured in complete medium containing 5 μg/ml puromycin for at least 14 days for infection-positive selection, until the proportion of fluorescent cells observed under a microscope reached 100%. The cells were then used for subsequent experiments.

### RNA-Seq

For RNA-Seq, biological duplicates of two independent Hela cell clones (total n=4) were seeded on 6-well dishes and grown for ∼48 hrs at 37°C to ∼75% confluence. Cells were then shifted to 32°C, 37°C and 40°C, and incubated for 24 hrs. For the okadaic acid treatment assay, HEK293T cells were seeded on 6-cm dishes in triplicates and grown overnight at 37°C. Cells were then shifted to 33°C or 39°C incubators for 12 hrs and then treated with okadaic acid (1μM) or DMSO as solvent control for 2 hrs at 33°C or 39°C, respectively. Total RNA from Hela and HEK293T was extracted as described below. Sequencing libraries were prepared by poly(A) selection using the TruSeq mRNA Library Preparation Kit. Sequencing was performed on an Illumina HiSeq 2500 system with V4 sequencing chemistry, generating around 50 million 150-bp paired-end reads per sample. RNA-Seq data are made publically available under GSE262498.

### Cloning

All cloning regarding RBM3 minigenes was done using the ClonExpress II One Step Cloning kit from Vazyme (C112). Briefly, the WT or mutated DNA fragments were generated by PCR, then the PCR fragments and the vector fragments from pcDNA3.1(+), which were digested with XhoI and HindIII enzyme, were incubated together for 30 min. Then, the reaction solution was directly added to DH5apha competent cells on ice for another 30 min, followed by heat shock of around 1 min at 42℃. Heat-shocked bacteria were immediately put on the ice for 5 min, followed by adding 300 μl of LB medium without antibiotics. These bacteria were then cultured at 37℃ for another 1hrs, and then they were plated on the agar plates containing ampicillin.

### Transfection

PEI (HY-K2014, MCE) was used as a carrier to transfect plasmids into cells. Briefly, around 1000 ng plasmid per well (12 well plate) was mixed with 100 μl of opti-MEM, and 3 μl of PEI (1μg/ml) was added into another tube containing 100 μl of opti-MEM. These two solutions were put at room temperature for ∼5 min, and then they were mixed at room temperature for another 20 mins. This mix was slowly added to overnight sub-cultured cells. WT or mutant mRBM3 and hRBM3 minigenes were transfected into cells at 37℃ overnight, then followed by different treatments if mentioned for another 24 hrs at different temperatures.

For siRNA (20 µM) transfection, RNATransMate (E607402, Sangon Biotech) served as the transfection reagent. In brief, approximately 10 µl of RNATransMate and 7 µl of siRNA per well (6-well plate) were diluted with 200 µl of serum-free DMEM medium, respectively. These two solutions were mixed and incubated at room temperature for 10 minutes to form the RNA/RNATransMate complex. Subsequently, this mixture was added to overnight-subcultured HT22 cells with approximately 60%-70% confluency. The cells were cultured for another 24 hrs to induce knockdown of the target gene, followed by subsequent chemical treatment.

### RNA extraction, RT-PCR and qPCR

For the HEK293T and N2a cells, Trizol (BS67.211.0100, Bio&SELL RNA Tri-Liquid) was used for RNA extraction as described in the user manual. Shortly, Trizol was directly added to cells after removal of the medium. Then, 1/5 volume CH_3_Cl was added and mixed, followed by 13000g centrifuge for around 30 min. The supernatant was added to the same volume of isopropanol, and then the mixture was centrifuged at 13000g for 30 min to pellet the RNA. The RNA pellet was washed with 70% ethanol (diluted with DEPC-treated water), and was dried at room temperature for 1min. The RNA was subjected to DNAse digestion at 37°C for at least 30 min, followed by PCI extraction and precipitation. Reverse transcription, RT-PCR and qPCR were performed as described previously^19^.

For the experiments of the hemin-induced HT22 cell model and the *in vivo* mouse model, brain tissues from the affected region (see *in vivo* mouse experiments) were ground with a grinder on ice and then dissolved in Trizol (R701-01, RNA-easy isolation reagent). For the HT22 cells treated in KCl, 4-AP and AFP treatment assay, Trizol (Invitrogen) was directly added into the cells after the culture medium was removed. Then, the cell lysates in Trizol were used for RNA extraction, and the procedures were the same as previously mentioned. Then, the cDNAs were generated as follows. For the qPCR of the hemin-induced HT22 cell model and the *in vivo* mouse model, 1st Strand cDNA Synthesis SuperMix for qPCR kit (11141ES60, YEASEN) was used to produce total cDNAs; for the HT22 cells transfected with minigenes in KCl, 4-AP and AFP treatment assay, minigene-specific transcripts were generated with the kit from Transgen (AT311, TransScript® One-Step gDNA Removal and cDNA Synthesis SuperMix) according to the manufacturer’s instructions with BGH-R primer. Briefly, around 500 ng of total RNA was incubated with the YEASEN or Tansgen Supermix containing reverse transcriptase, cDNA remover and RT-primer for 30 mins at 42 °C, followed by heating for 5 seconds at 85°C.

All the primers used are shown in the key resources table. For RT-PCR, the skipping and inclusion bands were amplified by corresponding gene-specific primers, and amplified bands were detected by agarose gels. The RT-PCR data in HEK293T and N2a cells was quantified with ImageQuant TL software. For the RT-PCR data in HT22, ImageJ/Fiji was used to quantify the data. For the qPCR experiments in HEK293T and N2a cells, mRNA was normalized using hHPRT and mHPRT, respectively. For the qPCR experiments in HT22 cells and *in vivo* mouse models, mRNA expression levels were normalized using mGAPDH for qPCR analyses.

### CD spectroscopy analysis

Briefly, the CD (circular dichroism) signal of rG4 was checked using a Jasco J-1500 CD spectrophotometer and a 0.1-cm path length quartz cuvette (Hellma Analytics) was employed using a volume of 200 μl. Samples with 5 μM RNA (final concentration) were prepared in 50 mM KCl or 0.1 mM KCl in 50mM Tris-HCl pH7.5. Each of the RNA samples was then thoroughly mixed and denatured by heating at 95 °C for 5 min and cooled to room temperature overnight for renaturation. The RNA samples were scanned from 220–310 nm at different temperatures and spectra were acquired every 1 nm. All spectra reported were an average of 3 scans with a response time of 0.5 s/nm. They were then normalized to molar residue ellipticity.

### NMR

All 1D ^1^H spectra were recorded on a Bruker Avance III spectrometer equipped with a ^1^H/^13^C/^15^N TCI cryoprobe. The sample temperature was adjusted with a MeOD-d4 sample according to Findeisen *et al*^82^. The RNA sample (Phosphate-GGGGGUGGAGGGCAGCGGCAGA) was synthesized by Microsynth on a 0.2 μmol scale giving a yield of 23.7 nmol. The RNA was dissolved in 500 µl of a 95% H_2_O/5% D_2_O (100 atom%D, ARMAR Chemicals, 1070-1X10ML) solution yielding a concentration of 47 μM. A standard 5 mm NMR tube (TA, ARMAR Chemicals) was used. 1D ^1^H spectra were recorded using a gradient 1-1 echo sequence^83^ with the application of two pulse field gradients of 1 ms and 15.9 G/cm similar to the gradient enhanced 1-1 echo HMQC^84^. Pulse train delays of 45 μs and 90 μs were used for the first and second pulse pair, respectively. Spectra were recorded with 8192 complex points, a spectral width of 22.0 ppm, typically 512 transients and a relaxation delay of 1.5 s. The irradiation frequency was adjusted to the remaining water signal. Spectra were processed using a quadratic sine shifted by 90°. Chemical shifts were referenced to DSS (2,2-Dimethyl-2-silapentane-5-sulfonate sodium salt) using an external commercial standard sample for testing water suppression containing 2 mM sucrose and 0.5 mM DSS (Bruker Biospin), measured at exactly the same temperatures.

### Protein extraction and western blot

For the HEK293T and N2a cells, whole-cell extracts (WCEs) were prepared with lysis buffer (20mM Tris (pH 8.0), 2% NP-40 (v/v), 0,01% sodium deoxycholate (w/v), 4mM EDTA and 200mM NaCl) supplemented with protease inhibitor mix (Aprotinin, Leupeptin, Vanadat and PMSF). For the HT22 cells and brain samples from mice, WCEs were prepared with IP lysis buffer (G2038, Servicebio) containing cocktail protease inhibitor (G2006, Servicebio) with sonication. Protein concentrations were determined using BCA assay (23225, Pierce™ BCA Protein Assay Kit, Thermofisher) according to the manufacturer’s instructions. SDS-PAGE and Western blotting followed standard procedures. Western blots were quantified using the ImageQuant TL software (HEK293T and N2a cells normalized with hnRNPL) and ImageJ/Fiji (HT22 cells and mice normalized with GAPDH). The following antibodies were used for Western blotting: hnRNPL (4D11, Santa Cruz), RBM3 (14363-1-AP, Proteintech), GAPDH (A19056, Abclonal) and NeuN (66836-1-Ig, Proteintech).

### Bioinformatics analysis

To reveal temperature-dependent changes in exon inclusion, 150 nt paired-end RNA-Seq samples from HEK293T and HeLa were aligned to the human hg38 genome, using STAR (v2.6.1a). Exon inclusion levels were then calculated using rMATS (v3.1.0) and further filtered using standard Python code. To obtain only high confidence splicing changes upon temperature treatments, we compared each temperature in biological replicates (triplicates for HEK293T (duplicates in DMSO treatment at 33℃ in HEK293T) and four replicates for Hela).

Cassette exons (Percent Spliced-In (PSI) > 0.05 and PSI < 0.95, and exon length > 20 bp) were classified into three groups, including cold-repressed exons (CRE), cold-induced exons (CIE), and not temperature-sensitive (NS), based on the changes of PSI (delta PSI) values at different temperatures. In detail, for HEK293T cells, CRE and CIE were defined as those exons with |delta PSI| ≥ 0.1 and BH-adjusted p value < 0.05 in the comparison of cells at 35℃ *vs* 39℃. In addition, CLK-dependent (only passed the thresholds in DMSO-treated cells) and independent (passed the threshold in both DMSO-treated and TG003-treated cells) exons were analyzed separately (Figure 1D and Figure S1C). For Hela, CRE and CIE were defined similarly in two comparisons, 32℃ *vs* 37℃, and 37℃ *vs* 40℃, respectively. However, the final NT exons were defined by the intersection of NT in these two comparisons (Figure 1E). The sequence flanking each cassette exon, and the upstream and downstream exons (Figure 1B) were extracted using bedtools^85^.We further predicted G4 in these sequences by searching motifs from a previous study^48^, and using G4Hunter^49^. The G4 motifs were searched in 50 bp sequences flanking splice sites. For G4Hunter analysis, we scanned 200 bp sequences flanking splice sites with a 25 bp window at each base to get a nucleotide-resolution score. All exons were used to examine the general correlation between G4 score and delta PSI. Hela rG4 sequencing (rG4-seq) data were downloaded from the Gene Expression Omnibus (GEO) with accession number GSE77282. The identified rG4 regions with a stop signal in cells cultured in the presence of K^+^ or PDS (K^+^ and PDS) were converted from hg19 to hg38 coordinates with UCSC LiftOver and used for the downstream analysis. We calculated the number of exons overlapped with rG4 peaks at each position from −200 to 200 nt for cassette exons and smoothed it with a 30 nt window. To compare the rG4-seq peak frequency across the three categories of exons (CRE, CIE and NT), the Reverse Transcriptase Stalling (RTS) values of the overlapped peaks were accumulated separately for each category. To obtain consistently temperature-regulated exons in HEK293T and Hela, we relaxed the thresholds to define CRE and CIE by requiring BH-adjusted p value < 0.05 and delta PSI > 0 (CIE) or < 0 (CRE).

For reproducing results/figures in the RNA thermometer study, the original codes are available in Github (https://github.com/christear/G4splicing). For the splice site strength prediction, we used MaxEntScan (http://hollywood.mit.edu/burgelab/maxent/Xmaxentscan_scoreseq.html) to get the 5’-splice site strength. For sequence alignment, we employed DNASTAR to generate the alignment results. The schematic figures in this paper were made by Biorender (https://www.biorender.com/).

### CCK-8 assay and dead/live staining

HT22 cell viability was measured using the cell counting kit-8 (CCK-8) according to the manufacturer’s instructions (HY-K0301, MCE). Briefly, 2000 cells were seeded in 96-well plates and allowed to grow overnight before treatment with the indicated concentrations of 4-AP for 24 hrs. After treatment, the cell culture medium was replaced with 100 μL fresh medium containing 10 μL CCK-8 solution and incubated for 1–3 hrs in a humidified incubator. The absorbance at 450 nm was measured with a microplate reader. For the dead/live cell assay, live cells were stained with LiveDye (a cell-permeable green fluorescent dye), and dead cells were stained with NucleiDye (a cell non-permeable red fluorescent dye). In the dead/live staining (KTA1001, Abbkine), after treatment with 4-AP for 24 hrs, HT22 cells were washed with PBS and then incubated in buffer solution with LiveDye and NucleiDye for 30 min darkness. Cells were washed again with PBS, and the fluorescence was observed under a fluorescence microscope.

### Immunostaining

After the cultured cells were cleaned with PBS, HT22 cells were fixed with formaldehyde for 10 min and incubated with 0.25% Triton-X 100 for 30 min. The sections were dewaxed, rehydrated, boiled in pH 6.0 citrate antigen retrieval solution (G1206, Servicebio) and incubated with 0.25% Triton-X 100 for 30 min. After blocking with 5% bovine serum albumin (BSA) for 1 h, HT22 cells and the sections were directly incubated with the primary antibodies (mouse anti-NeuN (66836-1-Ig, Proteintech) and rabbit anti-RBM3 (14363-1-AP, Proteintech)) at 4°C overnight. After washing with PBS, the cells were incubated with 5% BSA containing the Alexa 488-or 594-conjugated secondary antibodies. After washing again, the cells and sections were immersed in 4’, 6-diamidino-2-phenylindole (DAPI) for 30 min. Immunofluorescence images were acquired with a fluorescence microscope.

### *In vivo* mouse experiments

The Animal Center of Wuhan University provided 20–25g adult male C57BL6/J mice. They were housed in standard conditions of 22℃ and 50–60% relative humidity with a 12 hrs light/dark cycle and free access to food and water. Animal experiments were approved by the Institutional Animal Care and Use Committee (IACUC) of Wuhan University (IACUC issue No.: WDRM-20240302B). The SAH mouse model was performed by the intravascular perforation method^86^. Mice were anesthetized with isoflurane. The left internal carotid artery (ICA) was exposed in the median neck incision. A silicone-coated monofilament was inserted from the left external carotid artery (ECA) along the ICA to the bifurcation of the anterior cerebral artery (ACA) and middle cerebral artery (MCA). When resistance was encountered, continue to push forward by 1 mm to puncture the blood vessel. Then monofilament was removed and the ECA was ligated. The sham group underwent the same surgical procedure without an endovascular puncture. Mice were intraperitoneally injected with 4-aminopyridine (4-AP) (HY-B0604, MCE) immediately at the dosage of 1mg/kg^87,88^ after SAH and 20 hrs after SAH. Mice were stereotaxically microinjected with lentiviral particles (GeneChem Co., Ltd.) containing lentivirus overexpressing RBM3 (OE-RBM3) (GeneChem Co., Ltd., NM_001166410) or a negative control (OE-NC) (GeneChem Co., Ltd.) and lentivirus sh-RBM3 (GTTGATCATGCAGGAAAGTctcgagAGACTTTCCTGCATGATCAAC) or sh-NC (GTCTCCACGCGCAGTACATTTctcgagAAATGTACTGCGCGTGGAGAC) at a dosage of 2.0 μl (5 × 108 TU/mL) into the left cerebral cortex 7 days before SAH onset. The stereotaxic coordinates are as follows: point 1, anteroposterior 0.3 mm from the bregma, mediolateral 3 mm from the midline, and dorsoventral 4 mm from the skull; point 2, anteroposterior 0.8 mm from the bregma, mediolateral 3 mm from the midline, and dorsoventral 4 mm from the skull. The injection speed was 0.4 μL/min, and the needle was left in place for 5 min after the injection. For the Hematoxylin and eosin (HE) and Nissl staining, the mice were deeply anesthetized 24 hrs after SAH and transcardially perfused with cold PBS followed by 10 ml of 10% paraformaldehyde. Whole brains were collected, fixed in 10% paraformaldehyde for 24 hrs, and then sectioned into 10-μm-thick coronal slices. For the samples of qPCR and WB, the mice were sacrificed by neck dissection immediately after transcardiac perfusion with cold PBS, and the affected cerebral cortex on the ipsilateral side of the bleeding site was removed and placed in a tube and immediately stored at −80°C for WB or were frozen in liquid nitrogen for qPCR.

### Mouse behavior test

Neurological function was detected using the modified Garcia score and the rotarod test at 24 hrs after SAH in mice. The modified Garcia score included six standards: spontaneous movement, limb movement, forelimb extension, climbing, tapping one limb, and response to facial irritation, each scoring 0–3. The total score ranges from 3 to 18 (see the table below). In the rotarod test, briefly, mice were placed on a rotating drum with a speed accelerating from 5 to 30 rpm within 2 min. One day in advance, mice were trained on a rotating rod until a stable performance level was reached. After 24 hrs of SAH, the mice were tested with an initial speed of 5 rpm/min accelerated to a maximum of 30 rpm/min. The duration of animals on the accelerating rotarod was recorded. High neurological test scores indicated better neurological function. All neurobehavioral tests were performed blinded. The modified Garcia Score table with expanded descriptions for each component^68,89^:

| Component | Description | Score Range |
| --- | --- | --- |
| Spontaneous Activity | 0: No spontaneous activity; 1: Minimal spontaneous activity with no coordination or purposeful movement; 2: Moderate spontaneous activity with some coordination and purposeful movement; 3: Normal spontaneous activity, with full coordination and purposeful movement. | 0-3 |
| Symmetry in the Movement of Four Limbs | 0: No movement of four limbs; 1: Asymmetric movement or no movement in one or more limbs; 2: Slightly asymmetric movement in all four limbs; 3: Symmetric movement in all four limbs. | 0-3 |
| Climbing on an Incline | 0: Unable to climb on an inclined surface; 1: Partially successful attempt to climb on an inclined surface; 2: Successful climb on an inclined surface with some hesitation or difficulty; 3: Successful climb on an inclined surface without hesitation or difficulty. | 0-3 |
| Symmetry of the Forelimbs Outstretching | 0: No forelimbs outstretching; 1: Asymmetric forelimbs outstretching; 2: Slightly asymmetric forelimbs outstretching; 3: Symmetric forelimbs outstretching. | 0-3 |
| Touch Response | 0: No response to touch stimuli; 1: Minimal response or delayed response to touch stimuli; 2: Moderate response to touch stimuli; 3: Normal response to touch stimuli. | 0-3 |
| Proprioception Response | 0: No proprioception response; 1: Partial or delayed proprioception response; 2: Slight reduction in proprioception response; 3: Normal proprioception response. | 0-3 |

### TUNEL staining

To detect neuronal death, double immunostaining of NeuN (66836-1-Ig, Proteintech) and TUNEL was conducted by TUNEL Assay Kit (C1089, Beyotime) in coronal sections at 24 hrs after SAH according to the manufacturer’s instructions.

### Hematoxylin and eosin (HE) and Nissl staining

Brain sections were embedded in paraffin and cut into 10 μm-sections. The sections were dewaxed, rehydrated, and stained with haematoxylin and eosin (C0105, Beyotime) or Nissl stain solution (G1036, Servicebio) according to the instructions. Scoring system for spongiosis evaluation: 0: absent spongiosis (no intercellular edema or vesicles/bullae); 1: almost no spongiosis (almost no intercellular edema or vesicles/bullae); 2: mild spongiosis (minimal intercellular edema, few small vesicles/bullae); 3: moderate spongiosis (moderate intercellular edema, multiple vesicles/bullae); 4: Severe spongiosis (marked intercellular edema, numerous large vesicles/bullae). The number of neurons with normal morphology and intensity of Nissl substance was measured by Image J.

### Intracellular K^+^ measurement

After treatment with 4-AP, the culture medium was removed, and the cells were washed with PBS and lysed in deionized water with sonication. 24 hrs after SAH, the cerebral cortex was taken and fully ground with deionized water. The protein concentration of the homogenates was quantified by a BCA protein assay kit. The K^+^ concentration was measured by a potassium (K^+^) turbidimetric assay kit (E-BC-K279-M, Elabscience) according to the manufacturer’s protocol. In brief, 80 μL homogenate was mixed with an equal volume of commercial protein precipitant, and after being centrifuged at 1100 × g for 10 min, 50 μL supernatant was added to a 96-well plate, and then 200 μL chromogenic agent was added to the wells and mixed fully with the supernatant. After being incubated at room temperature for 5 min, the optical density at 450 nm was measured with a microplate reader. The K^+^ concentration was normalized with the protein concentration in the homogenate lysates.

### Statistical analysis

All quantifications were performed with Image J if not mentioned. Figures 7H, 7K and 7R, Figures S5E and 5H are presented as box and whisker plots with min to max showing all points of biological replicates. Figure 7G is presented as a violin plot showing all points of biological replicates. The others are presented as scatter dot plots (line at mean with SD). All individual data points are shown. Statistical significance was determined using GraphPad Prism v10, using a Student’s t-test if not mentioned. Statistical significance was accepted at P ≤ 0.05. In the figure legends, “ns” denotes P ≥ 0.05, * denotes P ≤ 0.05, ** denotes P ≤ 0.01, ***P ≤ 0.001 and **** denotes P ≤ 0.0001.

## Acknowledgements

We are grateful for the assistance provided by Antje Grünwald and Bobby Viet Draegert from Freie Universität Berlin in the wet lab. We also appreciate the discussion and some help from Mary Kennedy, Luiza Zuvanov and other lab members in Prof. Florian Heyd group at Freie Universität Berlin. The circular dichroism (CD) spectroscopy analysis was performed in the lab of Prof. Dr. Beate Koksch at Freie Universität Berlin with help from Zeinab Mahfouz. This study was supported by the National Natural Science Foundation of China (No. 81971870 and No. 82172173) and the Deutsche Forschungsgemeinschaft (grant HE 5398/4-2 to FH). The authors thank Peter Schmieder and Monika Berbaum (FMP Berlin) for providing NMR pulse sequences and Jana Sticht (FU Berlin) for NMR data set templates.

## Author contribution

F.H. M.Z. and M.L. led and supervised the study.

M.Z. conceptualized the study with B.Z., F.H., C.L., and M.P..

M.Z. and C.L. performed most of the wet lab experiments and data analysis.

B.Z. did the bioinformatics analysis of rG4.

M.P. performed the RNA-Seq analysis.

M.A. did NMR experiments.

M.S. supervised the NMR experiments.

D.L. performed some *in vivo* mouse experiments.

A.E. prepared the samples for RNA-Seq (Hela).

S.M. prepared the samples for RNA-Seq (okadaic treatment sample).

M.L., W.L., S.C., L.W., L.Z., Q.H., Q.L., Q.H., R.G. and L.Y. provided advice and insights for neuronal damage in cells and *in vivo* mouse experiments to mimic clinic scenario.

L.Z. helped to analyze HE staining.

T.H. contributed valuable insights and discussions to this project.

M.Z., B.Z. and F.H. wrote the manuscript with the help from C.L., M.P., M.S., Y.H., M.L., X.Q., A.C., T.W., Q.H., and X.G..

