## Supplementary figures and figure legens for "Stabilizing mammalian RNA thermometer confers neuroprotection in subarachnoid hemorrhage"

### Supplementary figures and figure legends

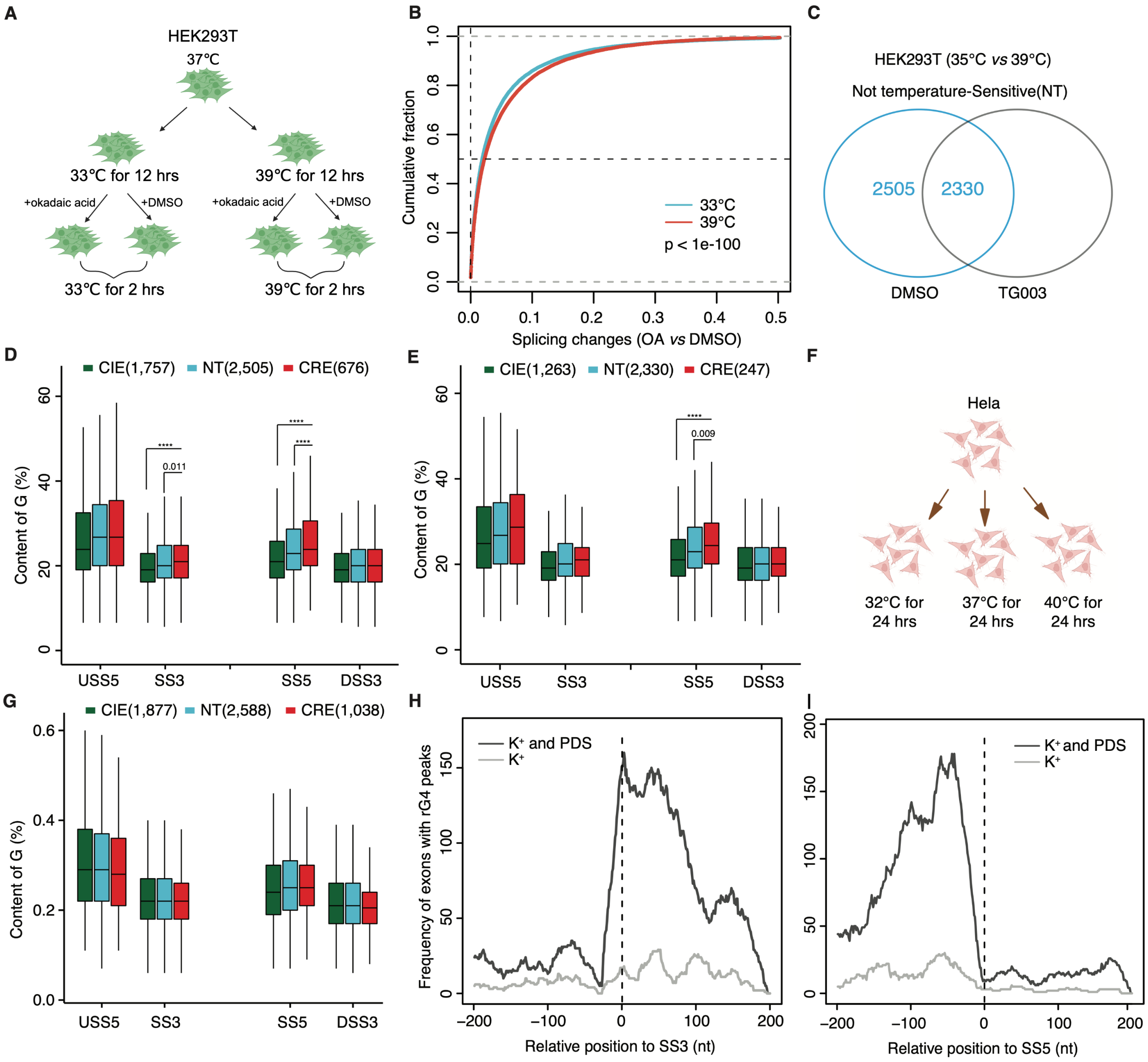


**Figure S1 G4 motifs are enriched around splice sites of cold-repressed exons (CRE), related to Figure 1.**

(A) Schematic illustrating HEK293T cells treated with okadaic acid (OA) and DMSO at 33°C and 39°C.

(B) Splicing changes upon OA treatment at high temperature (39°C) and low temperature (33°C) shown by the cumulative curves of absolute delta PSI values (OA *vs* DMSO) in HEK293T cells.

(C) Venn diagram showing the number of non-temperature-sensitive exons (NT) in HEK293T cells treated with DMSO and TG003 at 35°C *vs* 39°C (see Methods).

(D) and (E) Distribution of G-content in sequence within four regions (Figure 1C) of CRE, CIE and NT in HEK293T cells (35°C *vs* 39°C) that are dependent (D) or independent of CLK activity

(E) (see bioinformatics Methods). Significance was estimated by Wilcox test and only the significant results compared to CRE were indicated in the figure. ****, $p < 0.0001$.

(F) Schematic illustrating Hela cells cultured at 32°C, 37°C and 39°C.

(G) Distribution of G-content in sequence within four regions (Figure 1C) of CRE, CIE and NT in Hela cells (37°C *vs* 39°C).

(H) The frequency of exons with rG4 peaks around splice sites of cassette exons from rG4-seq data of Hela cells treated with $K^+$ and PDS or $K^+$ alone (see bioinformatics Method).

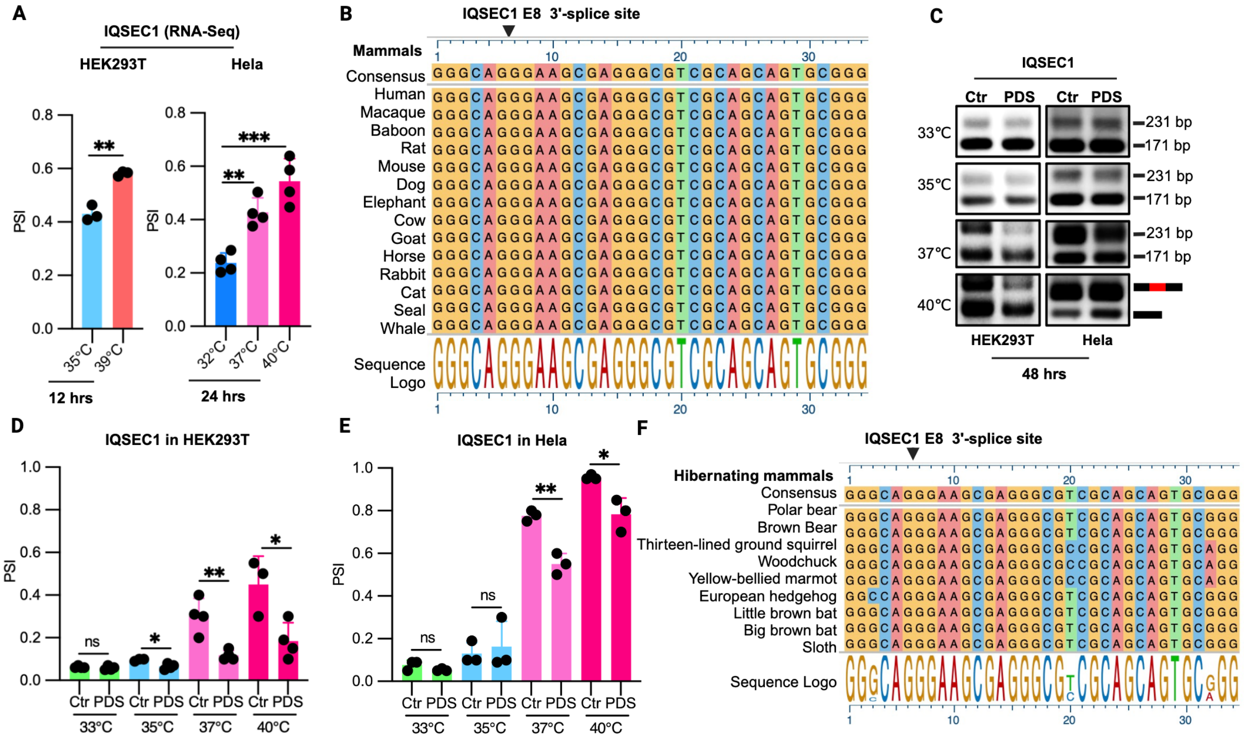


**Figure S2 G4 stabilizers reduce the inclusion of cold-repressed exons in IQSEC1, related to Figure 2.**

(A) Inclusion level (PSI) of the CRE E8 in IQSEC1 determined by RNA-seq analysis in HEK293T (left) and HeLa (right) cells (see Table 1). PSI values are based on rMATs.

(B) Evolutionary conservation of the G-rich element near the 3'-splice site of IQSEC exon 8 among different mammalian species.

(C)-(E) Inclusion level of IQSEC1 exon 8 after the G4 stabilizer (PDS) treatment shown by RT-PCR in HEK293T and Hela cells (C). HEK293T or Hela cells were treated with 10 μM PDS and cultured at 33°C, 35°C, 37°C, and 40°C for 48 hours, followed by RT-PCR. A representative gel image is shown. PCR products and sizes are indicated on the right. Representative gel images are shown in (C). Quantified data in (D) and (E) ($n \geq 3$).

(F) Evolutionary conservation of the G-rich element in the vicinity of IQSEC E8 in hibernating mammals (see bioinformatics Method).

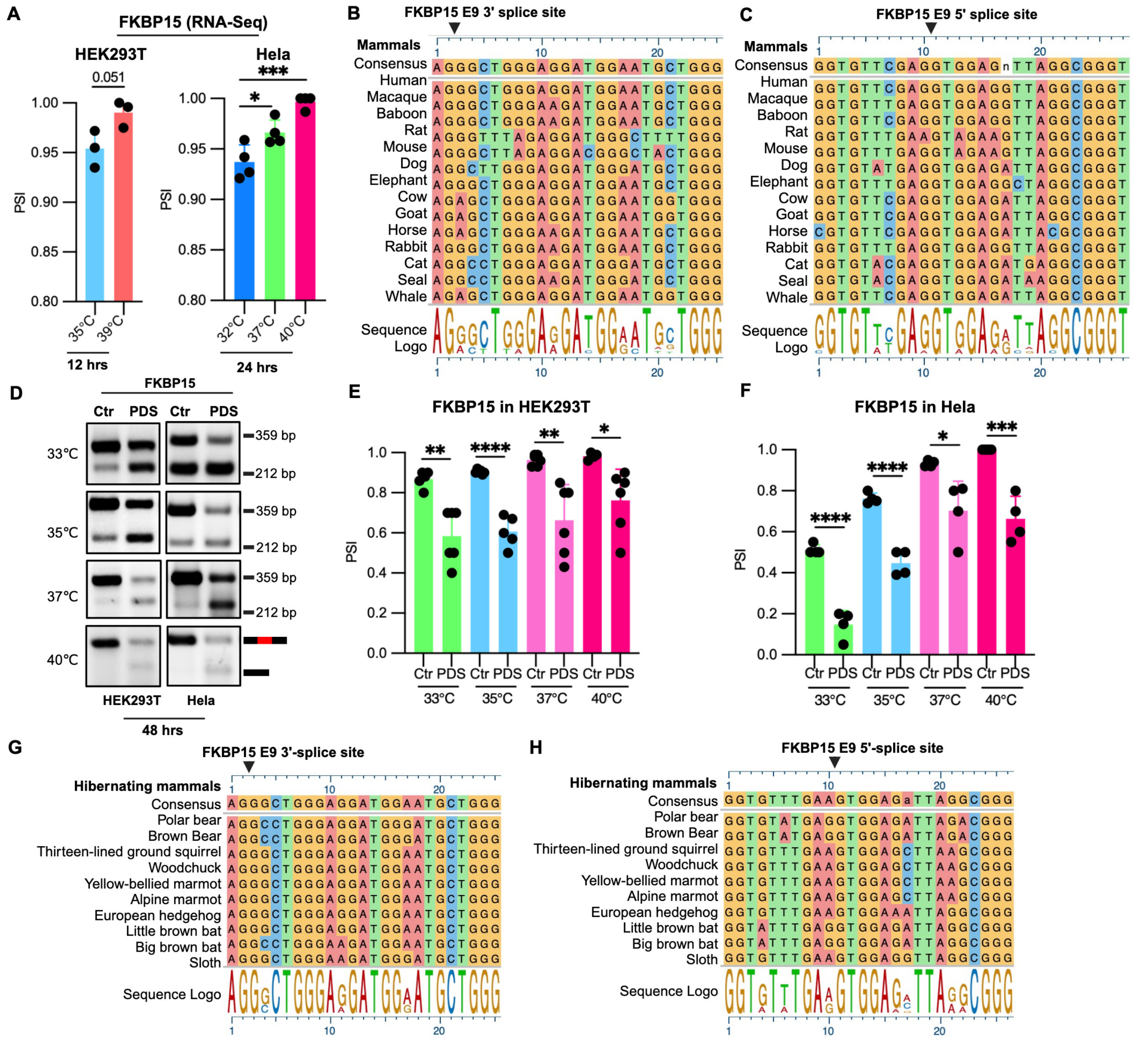


**Figure S3 G4 stabilizers reduce the inclusion of cold-repressed exons in FKBP15, related to Figure 2.**

(A) Inclusion level (PSI) of the CRE E9 in KBP15 determined by RNA-seq analysis in HEK293T (left) and HeLa (right) cells (See table 1). PSI values are based on rMATs.

(B) and (C) Evolutionarily conservation of the G-rich elements near the 3'-splice site (B) and 5'-splice site (C) of FKBP15 CRE E9 among different mammalian species.

(D)-(F) Inclusion level of FKBP15 CRE E9 after the G4 stabilizer (PDS) treatment shown by RT-PCR (D). Method as Figure S2C and Quantified data in (E) and (F) (n≥4).

(G) and (H) Evolutionary conservation of the G-rich elements in the vicinity of FKBP15 exon 9 in hibernating mammals.

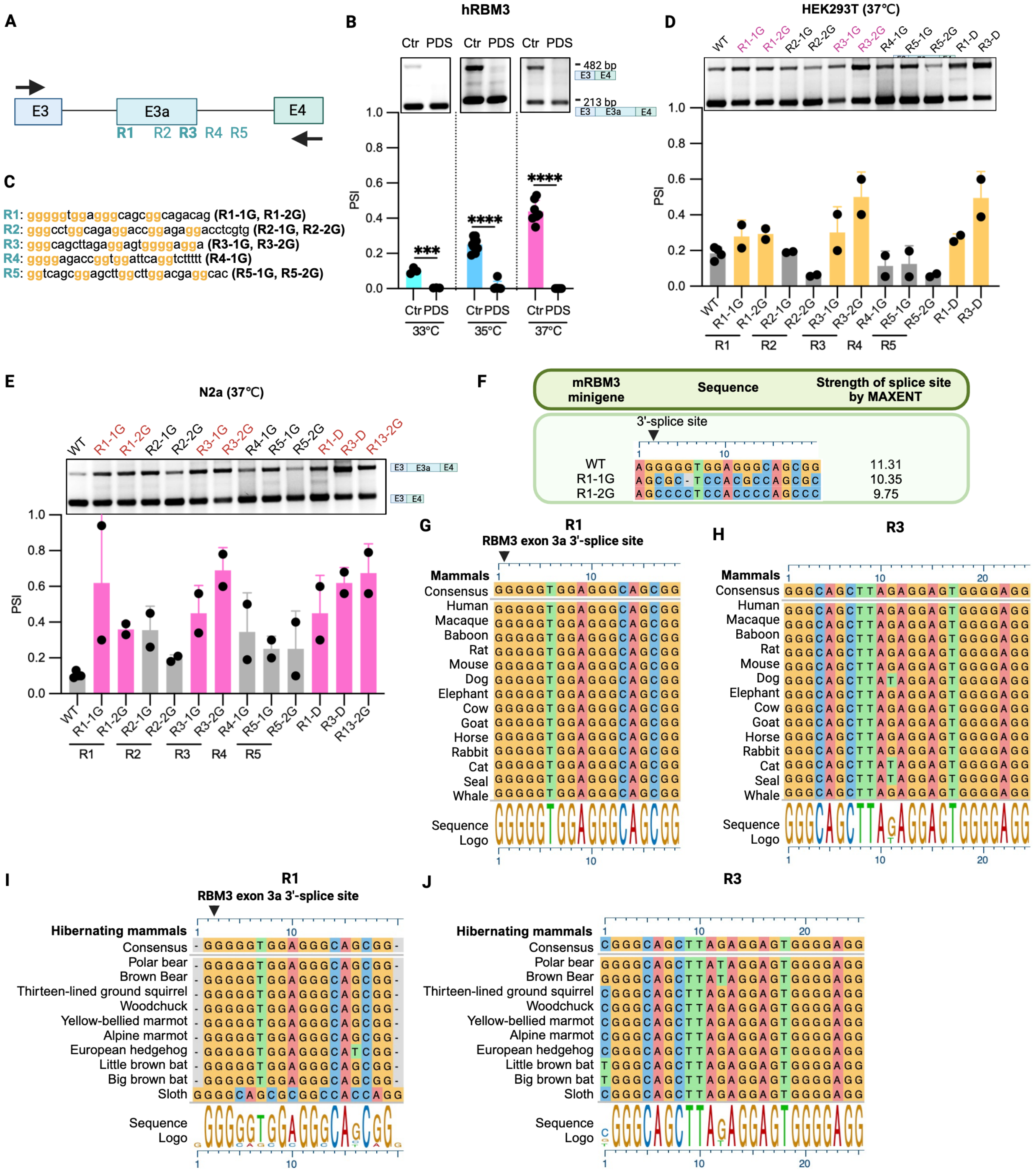


**Figure S4 Mutation of the conserved R1 and R3 elements decreases the inclusion of RBM3 exon 3a, related to Figure 3.**

(A) Illustration of positions of G-rich elements adjacent to the splice sites of RBM3 exon 3a. R1, R2, R3, R4 and R5 represent the position of each G-rich element, respectively.

(B) Inclusion levels of RBM3 exon 3a after treatment of HEK293T cells with the G4 stabilizer PDS in the hRBM3 minigene. Upper part depicts representative gels depict RT-PCR results, and the lower portion shows quantified results (see method, n=6 for 35°C, n=5 for 33°C and 37°C).

(C) WT and mutant sequences of G-rich elements in A. R1-1G, R2-1G, R3-1G, R4-1G and R5-1G represent one G to C mutation in every G4 tract, while R1-2G, R2-2G, R3-2G, R4-1G and R5-2G indicate the GG to CC mutation in every G4 tract.
(D) and (E) Inclusion levels of RBM3 exon 3a in WT and mutant mRBM3 minigene at 37°C in HEK293T (D) and N2a (E) cells (n=2, see Method).
(F) Splice site strength prediction for 3'-splice site of RBM3 exon 3a WT and mutants (see bioinformatics Method).
(G)-(J) Sequence alignment of R1 and R3 across multiple mammals (G) and (H), including hibernating species (I) and (J).

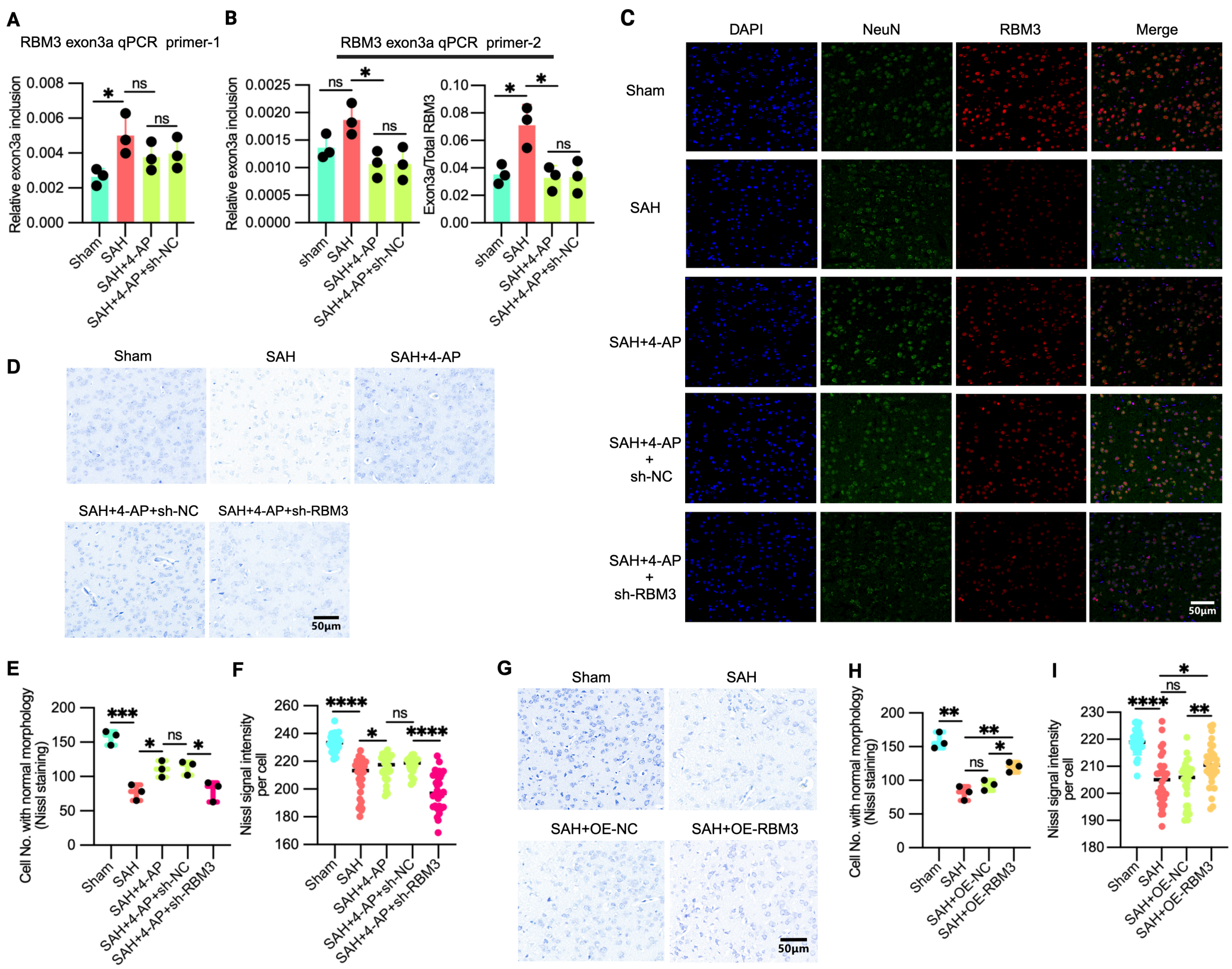


**Figure S5 4-AP administration protects from neuronal damage in a RBM3-dependent manner in the SAH mouse model, related to Figure 7.**

(A) and (B) RBM3 exon 3a inclusion and the ratio of RBM3 exon 3a to total RBM3 following 4-AP administration in the brain region (Figure 7B) of the SAH mouse model. RBM3 exon 3a qPCR primer-1 (A) and RBM3 exon 3a qPCR primer-2 (B) were used to quantify RBM3 exon 3a inclusion, respectively. Mouse GAPDH was used as normalized control (see Method, n=3 mice).

(C) Representative immunostaining images in the brain region (Figure 7B) of the SAH injected with lenti-shRBM3 and lenti-NC and sham mouse model *in vivo* after 4-AP and control administration (also see Method, n=3 mice).

(D) Representative images of Nissl staining of the mice treated as Figure 7D.

(E) Quantified number of Nissl substance-positive cells with normal morphology in (D) (n=3 mice).

(F) The intensity of Nissl substance, shown by Nissl staining in (D) (n=3 mice).

(G) Representative images of Nissl staining of the mice treated as Figure 7P (also see Figure 7A and Method, n=3 mice).

(H) Quantified number of Nissl substance-positive cells with normal morphology in (G) (n=3 mice).

(I) The intensity of Nissl substance, shown by Nissl staining in (G) (n=3 mice).
